# IRF8 regulates efficacy of therapeutic anti-CD20 monoclonal antibodies

**DOI:** 10.1101/2022.06.09.495444

**Authors:** Ludivine Grzelak, Ferdinand Roesch, Amaury Vaysse, Anne Biton, Françoise Porrot, Pierre-Henri Commère, Cyril Planchais, Hugo Mouquet, Marco Vignuzzi, Timothée Bruel, Olivier Schwartz

## Abstract

Anti-CD20 monoclonal antibodies such as Rituximab, Ofatumumab and Obinutuzumab are widely used to treat lymphomas and autoimmune diseases. They act by depleting B cells, mainly through Fc-dependent effectors functions. Some patients develop resistance to treatment but the underlying mechanisms are poorly understood. Here, we performed a genome-wide CRISPR/Cas9 screen to identify genes regulating the efficacy of anti-CD20 antibodies. We used as a model the killing of RAJI B cells by Rituximab through complement-dependent-cytotoxicity (CDC). As expected, the screen identified *MS4A1*, encoding CD20, the target of Rituximab. Among other identified genes, the role of Interferon Regulatory Factor 8 (IRF8) was validated in two B-cell lines. *IRF8* knockout also decreased the efficacy of antibody-dependent cellular cytotoxicity and phagocytosis (ADCC and ADCP) induced by anti-CD20 antibodies. We further show that IRF8 is necessary for efficient CD20 transcription. Levels of IRF8 and CD20 RNA or proteins correlated in normal B cells and in hundreds of malignant B cells. Therefore, IRF8 regulates CD20 expression and controls the depleting-capacity of anti-CD20 antibodies. Our results bring novel insights into the pathways underlying resistance to CD20-targeting immunotherapies.

## Introduction

CD20 is a B cell surface marker belonging to the MS4A (membrane-spanning 4-domain family A) protein family (1). B cells express CD20 from the late pre-B stage until plasmablast and plasma cell stages (2,3). Although CD20 function remains to be fully elucidated, the protein may act as a calcium ion channel and is involved in cell activation and differentiation (4–6).

Approximatively 90% of lymphomas are of B-cell origin (7). Non-Hodgkin lymphomas represent the sixth and seventh most common cancers in females and males, respectively, in the United States (8). The first anti-CD20 monoclonal antibody (mAb), Rituximab (RTX), has been approved in 1997 by the FDA to target B cells in non-Hodgkin lymphomas (9). RTX also demonstrated efficacy for the treatment of autoimmune diseases, such as rheumatoid arthritis and multiple sclerosis (10).

Additional anti-CD20 mAbs are in clinical use and have been classified into two types (type I and type II). Type I mAbs redistribute CD20 into lipid rafts, facilitating complement recruitment. Type II mAbs are less potent to elicit complement lysis but efficiently trigger direct cell death, because of increased cross-linking capacities (11). Type II mAbs binding to CD20 induces 1:2 (IgG:CD20) “terminal” complexes whereas type I mAbs form 1:2 or 2:1 “seeding” complexes able to recruit more mAbs or CD20 molecules (12). RTX and Ofatumumab (OFA) are of type I, whereas Obinutuzumab (OBZ) is a type II antibody. RTX is a chimeric murine/human glycosylated IgG1 with a human κ constant chain and murine light and heavy chains variable regions, OFA is of human origin and OBZ is a humanized antibody that harbors a modified Fc domain enhancing ADCC activity (13).

CD20-targeting immunotherapies are efficient to treat B-cell lymphomas, but resistances and relapses can occur (14,15). Resistance may be linked to low levels of CD20, “shaving” (i.e trogocytosis) of CD20-antibody complexes by macrophages (16) or CD20 internalization (17). Relapses can originate from subclones with low CD20 expression selected by the treatment (18). Consistently, high CD20 expression has been associated with a better survival in patients with B-cell lymphomas (19,20) and CD20 levels can decrease after Rituximab treatment (21). However, the mechanisms that control CD20 levels in malignant B cells and the CD20-low subclones are unknown.

The mechanism of action of anti-CD20 antibodies remains debated (22). Both direct induction of apoptosis and Fc-effector functions, such as antibody-dependent cellular cytotoxicity (ADCC), complement-dependent cytotoxicity (CDC) and antibody-dependent cellular phagocytosis (ADCP) may play a role. Depletion of B cells by ADCP and ADCC mediated by macrophages, monocytes or NK cells was demonstrated *in vitro* and in mice (23–26). Polymorphisms in FcγRIIIA in humans may impact the response to Rituximab (27–29). In clinical studies, RTX infusion induced complement consumption in patients with Chronic Lymphocytic Leukemia (CLL) (30). Furthermore, polymorphisms in C1q were associated with survival and RTX response in Follicular Lymphoma (FL) and Diffuse Large B-Cell Lymphoma (DLBCL) (31,32). In mice, anti-CD20 antibodies eliminate B cells by mediating their phagocytosis by liver resident macrophage Kupffer cells (33). Type I and II anti-CD20 also trigger cells death by caspase-dependent and independent mechanism, respectively (22,34,35). However, there is so far no direct evidence that this process is operative *in vivo* (36). The different mechanisms of action of anti-CD20 antibodies require sufficient CD20 surface levels on target cells.

CD20 expression is regulated by transcription factors such as PU.1, OCT1, OCT2 or NFκB (37–39) and by epigenetic mechanisms controlled by DNA methyltransferases and histones deacetylases (40,41). Whether other cellular proteins modulate the efficacy of anti-CD20 antibodies is not fully understood. Here, we sought to identify and characterize such proteins by performing a whole-genome CRISPR/Cas9 screening in RAJI cells subjected to CDC by RTX. We identified IRF8 as a protein regulating CD20 expression and modulating efficacy of therapeutic anti-CD20 antibodies in cell culture.’

## Results

### Identification of genes involved in the activity of anti-CD20 antibodies

We performed a whole genome CRISPR/Cas9 screen to identify genes involved in the sensitivity of B cells to anti-CD20-induced complement-dependent cytotoxicity (CDC). We used the genome-wide Toronto KnockOut (TKO) CRISPR Library v3, which consists of 71,090 guide RNAs (gRNA) targeting 18,053 genes (42). The library includes 4 gRNA per targeted gene and 142 irrelevant sequences targeting EGFP, LacZ and luciferase. Fig. 1A summarizes our screening approach. We transduced the RAJI B cell line, an EBV+ Burkitt lymphoma, with the TKOv3 library. We then submitted the transduced cells to complement-dependent lysis with Rituximab (RTX) and normal human serum (NHS). Surviving cells were either directly sorted (condition A) or cultured (condition B). High-throughput sequencing and Model-based Analysis of Genome-wide CRISPR-Cas9 Knockout (MAGeCK) enabled the identification of gRNAs enriched in surviving cells, using a false discovery rate (FDR) < 1%. MAGeCK takes into account all 4 gRNA for each gene to calculate an enrichment score (43). The main genes identified in condition A (Figure 1B, left panel) were *SMARCA4, AMBRA1, DHX29, CUL3, KHDC4, IRF8* and *MS4A1* (the gene coding for CD20). Genes found in condition B were *AMBRA1, CD19, KEAP1, MAPKAP1, CUL3, RING1, KHDC4, RICTOR, IRF8* and *MS4A1* (Figure 1B, right panel). In both conditions, gRNA targeting *MS4A1*, the gene coding for CD20, exhibited the highest level of enrichment, validating our screening approach (Figure 1B). The representation of gRNA was higher in condition A (94.5%) than in B (68.5%) (Figure 1C), consistent with the stronger level of selection applied in condition B. We considered as hits the genes identified in both conditions: *IRF8* (Interferon regulatory factor 8), *MS4A1* (Membrane-Spanning 4-Domains, Subfamily A, Member 1), *CUL3* (Cullin 3), *AMBRA1* (Activating Molecule in Beclin-1-Regulated Autophagy) and *KHDC4* (KH Domain Containing 4).

**Figure 1.**
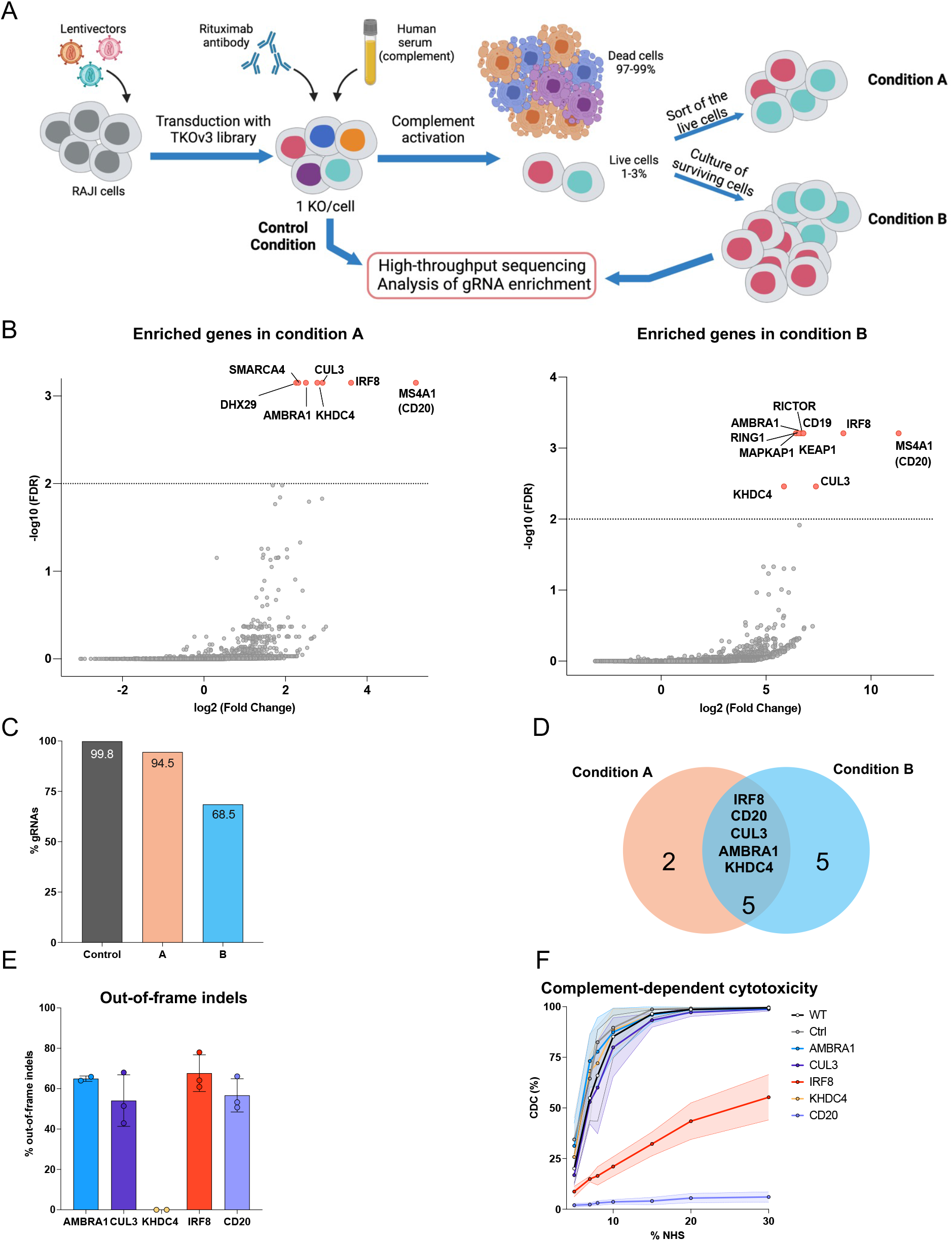
Top genes involved in complement-dependent cytotoxicity induced by anti-CD20 antibody rituximab (RTX) in a whole genome CRISPR-Cas9 screen. **A**. Protocol used for identification of genes involved in CDC. Two conditions were performed: in condition A, a sort of remaining live cells was performed 24 hours post complement activation while in condition B the surviving cells were cultured for 8 days before DNA extraction. **B**. Visualization of the enrichment of gRNA-coding sequences in conditions A (left panel) or B (right panel) represented with minus logarithm base 10 of the false discovery rate (FDR) on y-axis and logarithm base 2 of the fold change compared to control condition on x-axis. Genes with a FDR<1% are identified on the graph. **C**. Percentage of gRNA-coding sequences detected in each condition: control, A or B on the total number of gRNA in the library TKOv3. **D**. Venn diagram representing the number of genes enriched in conditions A, B or both. **E**. Percentage of out-of-frame indels for each gRNA used to knock-out cells. IRF8 KO cells were electroporated with a mix of 3 gRNAs and the percentage depicted is the maximum of the 3. **F**. Percentage of CDC in RAJI knocked-out for the top genes enriched in the screen with the indicated concentrations of normal human serum (NHS) added as a source of complement. The mean of 3 experiments is represented. The dotted line is the standard deviation and the area between the curve and the standard deviation is shaded. To take into account direct apoptosis triggered by RTX, the percentage of CDC was calculated as: (% dead cells in condition - % dead cells in no serum condition)/(100-% dead cells in no serum condition).

To validate these results, we engineered cells with individual CRISPR-Cas9 knockouts (KO) for each of the five hits. We used as controls wild-type RAJI cells (WT) and cells electroporated with a control gRNA (Ctrl). To assess the levels of gene editing (and ultimately of KO) in these pooled cell populations, we used the Tracking of Indels by Decomposition (TIDE) method (44), which measures the percentage of out-of-frame indels (Figure 1E). In these conditions, *AMBRA1, CUL3, IRF8* and *MS4A1* were efficiently suppressed, with around 60% of out-of-frame indels, but *KHDC4* was not. We next evaluated the RTX-mediated CDC sensitivity of each of KO cells, in the presence of increasing amounts of serum as a source of complement (Figure 1F). WT and Ctrl cells were sensitive to CDC, with nearly 100% killing at high serum concentration. *MS4A1* KO cells displayed near-complete protection against killing. *IRF8* KO cells significantly resisted lysis. None of the other hits (*AMBRA1, CUL3, KHDC4)* impaired the activity of RTX (Figure 1F). Altogether, these results indicate that MS4A1 and IRF8 genes regulate RTX-mediated CDC of RAJI cells.

### IRF8 controls CD20 expression and RTX-induced CDC

We next investigated the interplay between IRF8 and CD20. We first assessed *IRF8* KO efficiency by using RT-qPCR and flow cytometry (Figure 2A). In *IRF8* KO cells, *IRF8* mRNA levels were significantly decreased compared to control cells (p<0.005, relative expression of 0.43(±0.099 (SD)) *vs* 4.0 (±1.07)). IRF8 was diminished at the protein level by about 3.8-fold in nMFI (normalized Median Fluorescence Intensity) measured by flow cytometry (Figure 2A). There was no impact of *CD20* KO on either mRNA or protein levels of IRF8. Then, we rescued the expression of IRF8 in IRF8-KO cells by transduction with a lentivector expressing *IRF8*-*GFP* and subsequent sorting of GFP^+^ cells. Rescue of IRF8 expression was partial but significant with a 2.7-fold increase in nMFI (0.71±0.22 compared to 0.26, p-value = 0.0002; Mann-Whitney test). The rescue experiment indicated that that the decrease of sensitivity to CDC was due to CD20 down-regulation and not to an off-target effect of the CRISPR-Cas9 strategy. Of note, KO of *IRF8* did not impact the expression of complement-regulatory proteins CD46, CD55, CD59 and CR1. Some proteins were up-regulated (such as CD86) or down-regulated (such as CD21) in *IRF8* KO cells (Figure S3). This transcription factor may thus control the expression of a various proteins in B cell lines. We then tested the susceptibility of KO cells to CDC induced by RTX with 2 readouts: flow cytometry (Figure 2B, left panel) and high-throughput live imaging (Figure 2B, middle and right panel). Both approaches showed reduced mortality in *IRF8* KO cells (24.3 ± 11.6% for left panel) compared to control cells (73.3±6.8%), whereas the *IRF8* KO+IRF8 cells were not statistically different from the controls (56.1±10% CDC). The protection brought by *IRF8* KO did not however reach the extent of *MS4A1* KO (10.2±6% CDC). We also tested CDC triggered by an irrelevant anti-HLA-I antibody to assess the specificity of the protection. There was no significant effect of *IRF8* KO in the killing mediated by anti-HLA-I antibodies (Figure S2). Therefore, the reduced susceptibility of these cells to killing by RTX is due to CD20 down-regulation and not to the lack of IRF8.

**Figure 2.**
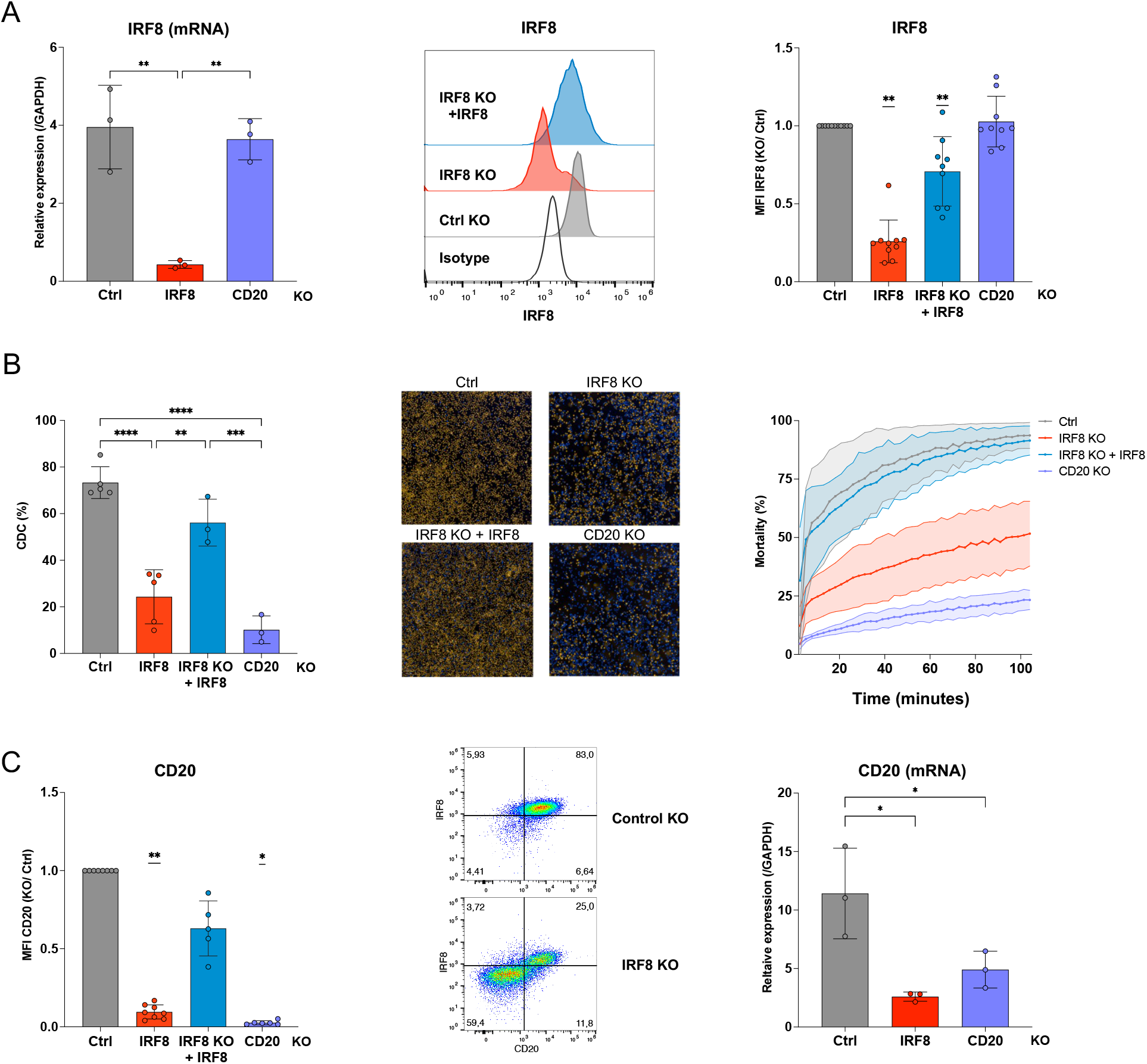
Effect of IRF8 KO in RAJI cells. **A. (Left panel)** mRNA levels of IRF8 in control, IRF8 KO or CD20 KO cells. mRNA levels of GAPDH were used to standardize the samples. A Shapiro-Wilk test confirmed normality of the data. Differences in expression were assessed by a one-way ANOVA test, **:p-value <0.005. **(Middle panel)** Representative histogram of flow cytometry staining for IRF8 protein. **(Right panel)** Normalized MFI of IRF8 protein (KO/Ctrl) for n=9-10 independent experiments, each point corresponds to one experiment. As normality was not confirmed, a Wilcoxon signed rank test was performed, **: p-value<0.01. **B**. CDC of RAJI KO analyzed by flow cytometry (left panel) or imaged with a high-throughput microscope (middle and right panels). (**Left panel**) Percentage of CDC in RAJI cells Control, IRF8 KO, CD20 KO or IRF8 KO complemented with IRF8, n=3-5. An ordinary one-way ANOVA was performed, **: p-value<0.005, ***: p<0.001, ****: p<0.0001. **(Middle panel)** Image of RAJI KO cells after 3h28 of complement and antibody exposure. Dead cells are depicted in orange (propidium iodide staining) and nuclei are in blue (DAPI staining). **(Right panel)** Percentage of mortality calculated as the number of dead cells/ number of nuclei depending on time. Standard deviations are depicted as shaded area. **C**. CD20 expression in RAJI-KO cells. **(Left panel)** Normalized MFI of CD20 protein (KO/Ctrl) for n= 5-8 independent experiments. A Wilcoxon signed rank test was performed, *: p<0.05, **: p-value<0.01. **(Middle panel)** Representative dot plots of CD20 and IRF8 staining in control or IRF8 KO cells. **(Right panel)** Relative expression of CD20 normalized to GAPDH. A one-way ANOVA test was performed, *:p-value<0.05.

We further examined the effect of knocking out *IRF8* on CD20 levels. The nMFI of CD20 in *IRF8* KO cells was strongly reduced, with a mean of 0.1±0.05 compared to 0.03±0.01 for *MS4A1* KO. The rescue of IRF8 expression increased CD20 levels by 6-fold (nMFI of 0.6±0.2 Figure 2C, left panel). A double staining of IRF8 and CD20 showed that control cells were positive for both proteins (Figure 2C, middle panel). Conversely, in the *IRF8* KO population, the fraction of IRF8 negative cells was also CD20 negative. As IRF8 is a transcription factor, we next asked whether the absence of IRF8 reduces the levels of CD20 mRNA. To this aim, we measured by RTqPCR (Figure 2C, right panel) the amounts of CD20 mRNA in the different KO cells. CD20 mRNA levels were 4.4 times lower in *IRF8* KO than in control cells (2.6±0.36 for *IRF8* KO and 11.41±3.87 for Ctrl). This decrease was similar to that in *MS4A1* KO (4.9±1.57). Therefore, IRF8 is necessary for efficient CD20 transcription in RAJI cells. Its absence induces resistance of these cells to RTX-mediated CDC.

### IRF8 drives CD20 expression in B cell lines

We then asked whether IRF8 controls CD20 expression in other B lymphoma cell lines. We first analyzed CD20 and IRF8 expression in SU-DHL-4 cells, an EBV-negative B cell line isolated from a DLBCL of the germinal center B-cell (GCB) type. We silenced *IRF8* in these cells (Figure S1). In *IRF8* KO SU-DHL-4 cells, *IRF8* mRNA and protein levels were decreased by 4.6-fold (p=0.005,2.0±0.7 *versus* 9.5±2.5) and 6.7-fold (0.15±0.018, significantly different from 1, p<0.0001) respectively (Figure 3A). RTX-induced CDC in *IRF8* KO cells was decreased by up to 6.7-fold (p<0.005) at different concentrations of serum (Figure 3B). We observed a 2.6-fold decrease in CD20 protein levels in *IRF8* KO cells (nMFI= 0.38±0.05, p<0.05) and a 49-fold decrease (p<0.05) in *MS4A1* KO cells (Figure 3C, left and middle panels). CD20 mRNA levels were also 3.3-fold lower in *IRF8* KO than in control cells (13.6±4.6 *versus* 45.1±12.2, p<0.01), which is similar to the decrease in *MS4A1* KO cells (20.1±4.4, p <0.05). Therefore, IRF8 controls CD20 transcription and susceptibility to RTX in SU-DHL-4 cells.

**Figure 3.**
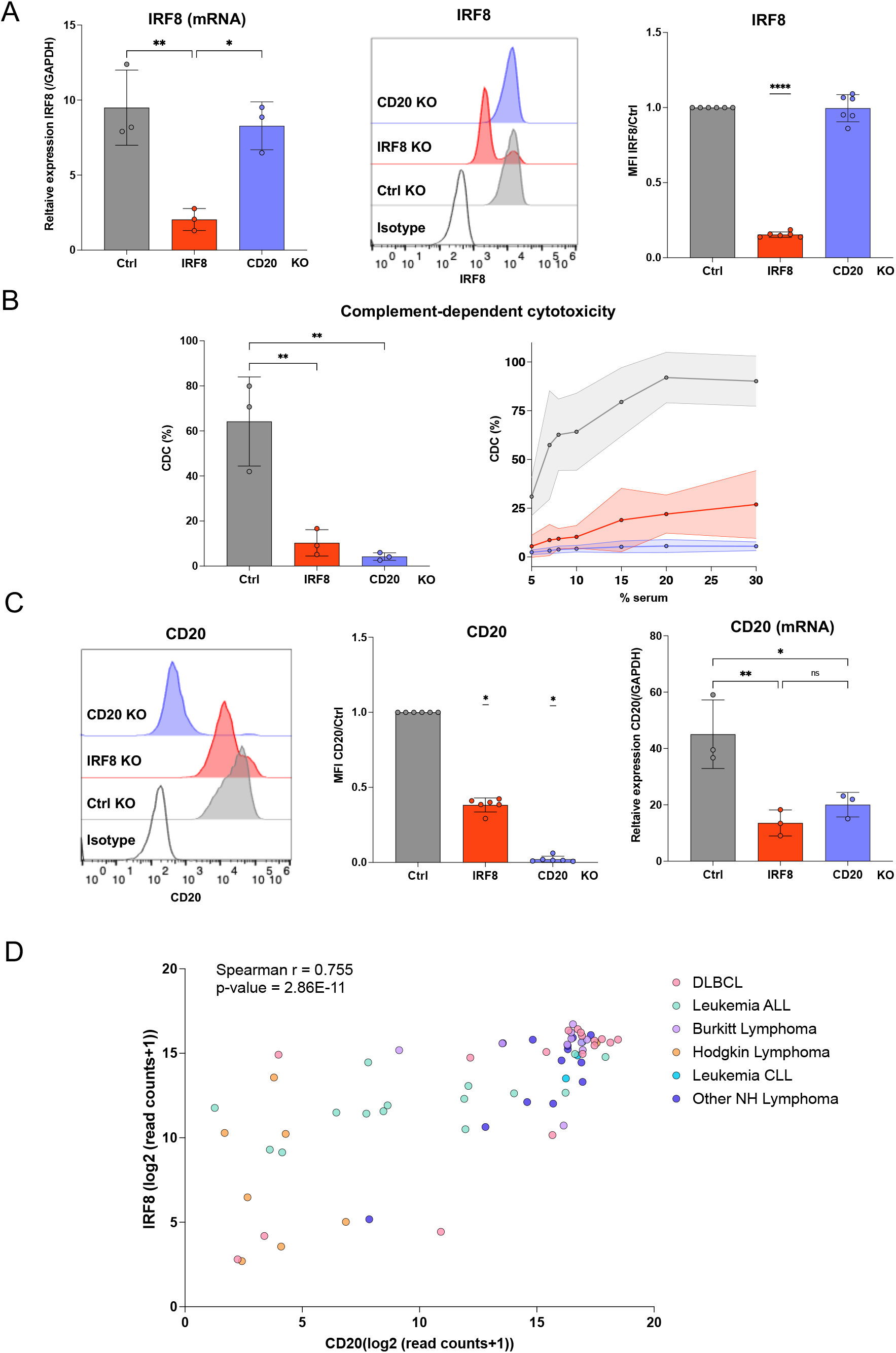
Effect of IRF8 in SU-DHL-4 cells and analysis of IRF8 and CD20 levels in a panel of B cell lines. **A. (Left panel)** Relative expression of IRF8 in control, IRF8 KO or CD20 KO cells normalized to GAPDH. A Shapiro-Wilk test confirmed normality of the data and differences in expression were assessed by a one-way ANOVA test, *: p<0.05, **: p-value = 0.005. **(Middle panel)** Representative histogram of flow cytometry staining for IRF8 protein. **(Right panel)** Normalized MFI of IRF8 protein (KO/Ctrl) for n=6 independent experiments, each point corresponds to one experiment. As normality was confirmed, a one sample t-test was performed, ****: p-value<0.0001. **B**. CDC of SU-DHL-4 KO analyzed by flow cytometry after one hour. **(Left panel)** Percentage of CDC in Control, IRF8 KO or CD20 KO cells with 10% normal human serum after 24h, n=3 independent experiments with independent knockouts, 6 days post-electroporation. An ordinary one-way ANOVA was performed, **: p-value<0.005. **(Right panel)** Percentage of CDC depending on the percentage of serum added, n= 3 independent experiments and KO. **C**. CD20 expression in SU-DHL-4 KO cells. **(Left panel)** Representative histograms of CD20 staining. (**Middle panel)** Normalized MFI of CD20 protein (KO/Ctrl) for n= 6 independent experiments. A Wilcoxon signed rank test was performed, *: p<0.05. **(Right panel)** Relative expression of CD20 normalized on GAPDH. A one-way ANOVA test was performed, *: p-value<0.05, **: p<0.01. **D**. Expression of IRF8 and CD20 represented in logarithm 2 of the read counts+1 in 66 B cell lines. ALL: Acute Lymphocytic Leukemia, CLL: Chronic Lymphocytic Leukemia, DLBCL: Diffuse Large B-Cell Lymphoma, NH: non-Hodgkin lymphoma.

To extend our observations, we assessed the correlation between *MS4A1* and *IRF8* mRNA levels using RNA sequencing data from the Cancer Cell Line Encyclopedia (CCLE) (Figure 3D). We investigated 66 B cell lines originating from Acute Lymphocytic Leukemia (ALL), Chronic Lymphocytic Leukemia (CLL), Burkitt lymphoma, DLBCLs, Hodgkin lymphoma and other non-Hodgkin lymphomas. There was a significant correlation between *IRF8* and *MS4A1* expression in these cells (Spearman coefficient of 0.755 and a p-value of 2.86·10^−11^) (Figure 3D). There were significant correlations for cells from DLBCLs, acute lymphocytic leukemia and other non-Hodgkin lymphomas but not from Burkitt and Hodgkin lymphomas (Supplementary Figure S4). Together, these results strongly suggest that IRF8 plays a role in the regulation of CD20 in various B-cell cancer models.

### Assessment of IRF8 and CD20 expression in primary B cell subsets

To determine whether IRF8 controls CD20 levels in non-malignant B cells, we examined IRF8 and CD20 expression in peripheral blood B cell subsets isolated from healthy donors (Figure 4). The gating strategy to identify B cells (expressing CD19) is detailed in Figure S5. Briefly, we stained cells with antibodies against CD19, CD27, CD38 and CD21 to identify different subsets: tissue-like memory B cells (CD19^+^ CD38^-^ CD27^-^ CD21^-^), activated-memory B cells (CD19^+^ CD38^-^ CD27^+^ CD21^-^), resting memory B cells (CD19^+^ CD38^-^ CD27^+^ CD21^+^), mature naïve B cells (CD19^+^ CD38^-^ CD27^-^ CD21^+^), plasmocytes/plasmablasts (CD19^+^ CD38^high^ CD27^high^), activated germinal center B cells (CD19^+^ CD38^+^ CD27^+^) and immature/transitional B cells (CD19^+^ CD38^+^ CD27^-^ CD21^+^). These stainings show that most B cell subtypes express both CD20 and IRF8. A subpopulation of tissue-like memory and activated memory B cells expressed IRF8 but not CD20. Thus, all CD20^+^ circulating B cells express IRF8, while not all IRF8 positive cells express CD20. This observation suggests that IRF8 is necessary, but not sufficient for CD20 expression in healthy donors-derived B cells.

**Figure 4.**
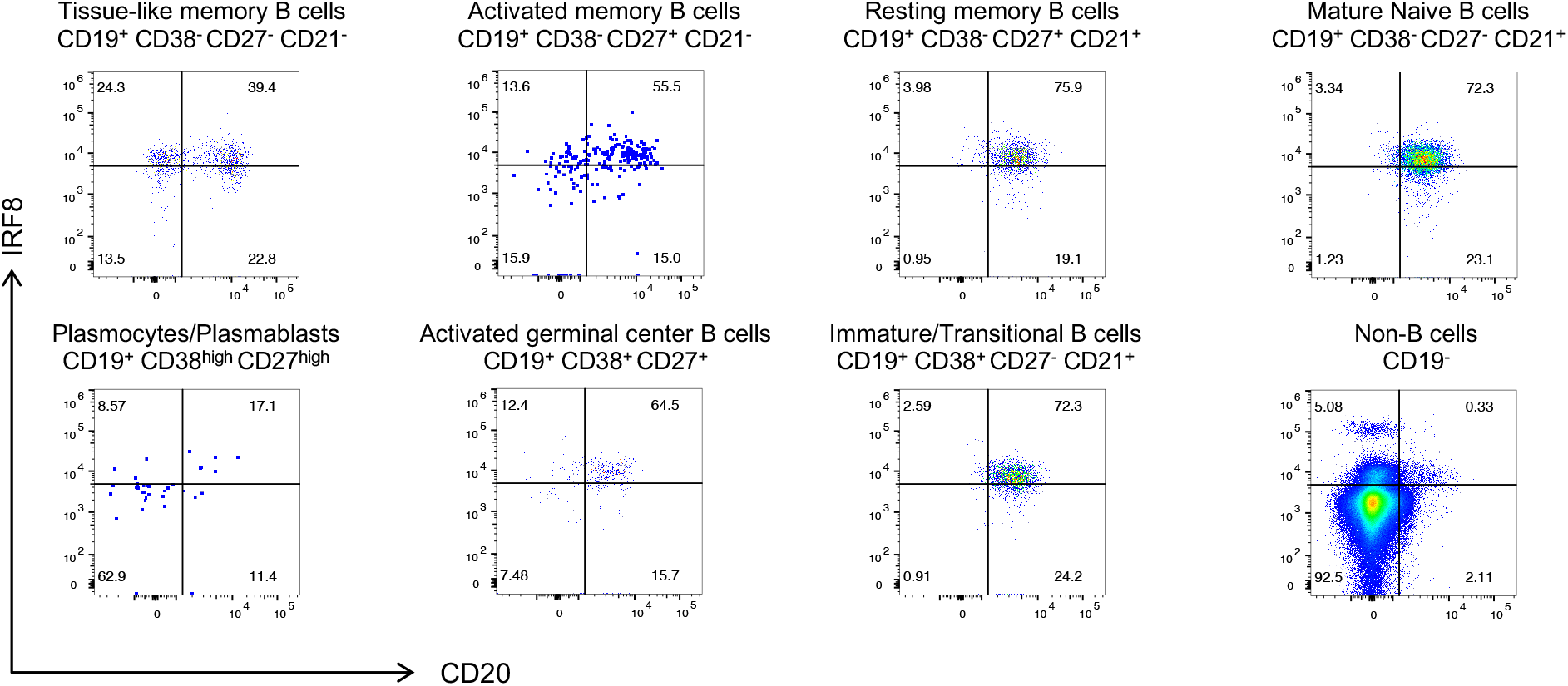
CD20 and IRF8 expression in primary B cells at various differentiation stages. **A**. IRF8 and CD20 staining in tissue-like memory, activated memory, resting memory, mature naive, activated germinal center, immature/transitional B cells, plasmocytes/plasmablasts and non-B cells. Staining of 1 donor representative of 4, performed in 2 independent experiments. The other donors are shown in supplementary figure 4.

### IRF8 impacts anti-CD20 activities of RTX, OFA and OBZ

We then investigated whether IRF8 regulates the activity of two other therapeutic anti-CD20 antibodies, OFA and OBZ. We assessed their activity in the CDC assay as well as in two surrogate assays for antibody-dependent cell cytotoxicity (ADCC) and antibody-dependent cell phagocytosis (ADCP).

The ADCC reporter test is based on Jurkat cells expressing CD16 (FcγRIII) and an NFAT response element driving expression of firefly luciferase. Addition of anti-CD20 antibodies on RAJI cells will bridge RAJI and Jurkat cells, activate the NFAT pathway and trigger a luciferase signal in Jurkat cells (45). The surrogate ADCP test is a flow-cytometry based assay using THP-1 cells as effector cells and RAJI cells as targets. The assay measures the triggering of RAJI/THP1 doublets by the antibodies and the subsequent phagocytosis of RAJI cells by THP-1 cells (46).

We first tested the binding of each antibody on RAJI *IRF8* and *MS4A1* KO cells. The MFI on control cells was similar for RTX, OFA and OBZ (42,015±13,408, 46,709±15,418 and 33,898±12,308 respectively) strongly suggesting similar affinity and avidity. *IRF8* KO decreased binding of the three antibodies by 15-, 13- and 15-fold for RTX, OFA and OBZ respectively (Figure 5A). *MS4A1* KO cells also displayed low antibody binding. *IRF8* KO cells were poorly sensitive to CDC (Figure 5B). At the highest antibody concentration (30 μg/ml) killing was decreased by a 4.4-, 4.1- and 7.7-fold for RTX, OFA and OBZ. Measurement of areas under the curves confirmed this decrease (Figure S6D). *MS4A1* KO cells were resistant to CDC with the three antibodies.

**Figure 5.**
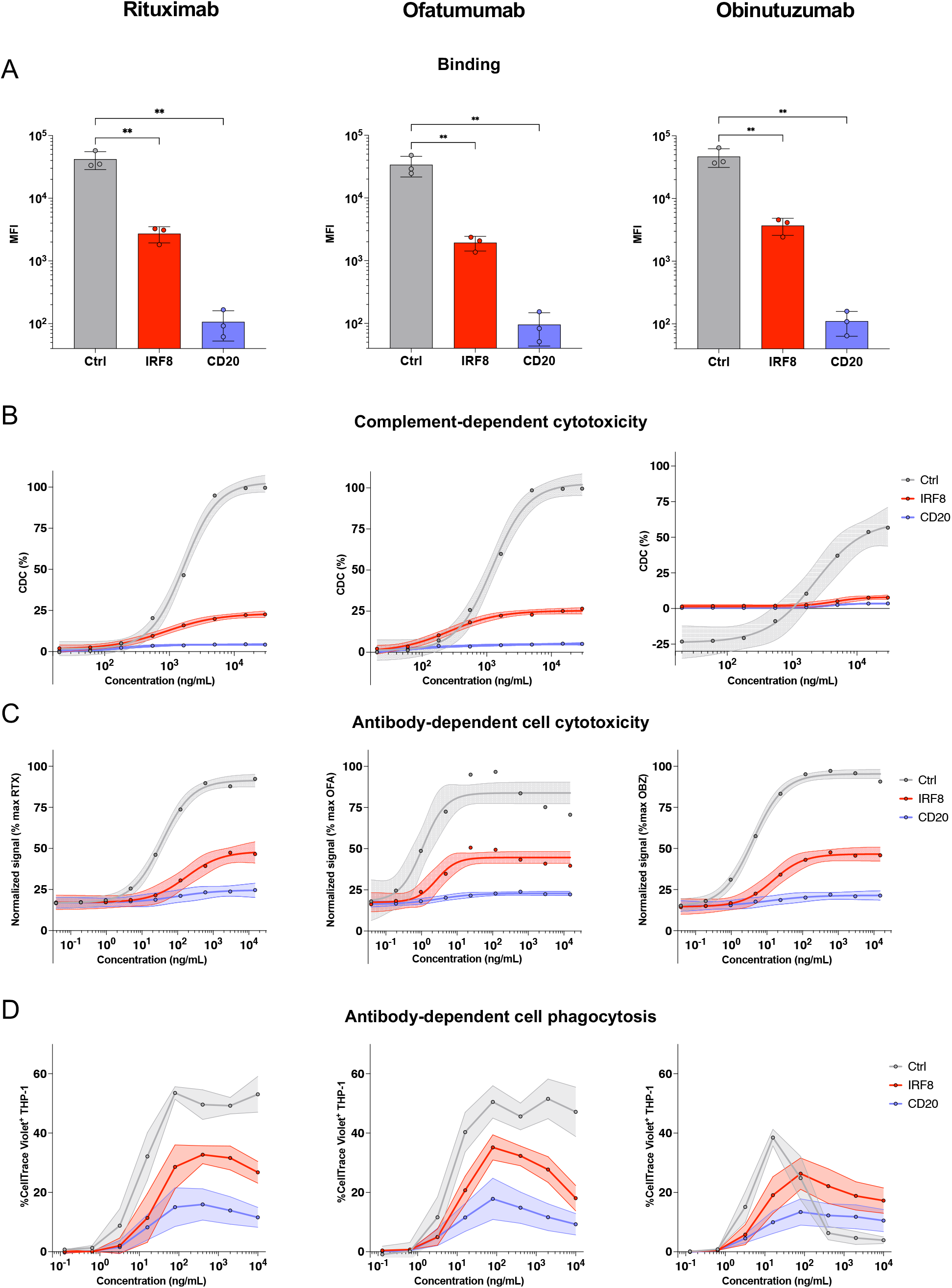
Effector functions of 3 anti-CD20 antibodies: Rituximab (left panels), Ofatumumab (middle panels) and Obinutuzumab (right panels). **A**. Antibody binding on RAJI KO cells. Median fluorescence of intensity is represented for n=3 independent experiments. Normal distribution was checked by a Shapiro-Wilk test which enabled an ordinary one-way ANOVA test to be performed. **: p-value <0.01. **B**. Dose response analysis of CDC induced by each anti-CD20 antibody. Concentrations are in ng/mL. Control KO, IRF8 KO and CD20 KO RAJI cells were used as targets. Results are from 3 independent experiments. **C**. Dose response of the surrogate antibody-dependent cell cytotoxicity test (ADCC) induced by each anti-CD20 antibody. Concentrations are in ng/mL, n= 3 independent experiments. For B and C non-linear sigmoidal 4 parameter logistic regressions were calculated and plotted on the graph. The 95% confidence interval of the regression is represented with the shaded area. Effective concentrations 50 and areas under the curve are shown in supplementary Figure 4. **D**. Dose response of the surrogate antibody-dependent cell phagocytosis test (ADCP) induced by each anti-CD20 antibody. Concentrations are in ng/mL. The mean of 4 independent experiments is represented with the standard deviation in shaded area. Non-linear sigmoidal 4 parameter logistic regressions were also performed but not represented due to the Hook effect observed for Obinutuzumab. To take this effect into account the regression was calculated for concentrations under 16 ng/mL. Effective concentrations 50 and areas under the curve are shown in supplementary figure 5.

*IRF8* KO cells were partially protected from ADCC, to a lesser extent than CDC, with a reduction of signal of 2.1-, 2.2- and 1.9-fold, with the three antibodies, respectively (Figure 5C). The variations between CDC and ADCC assays may reflect the different sensitivities of the readout (cell viability or luciferase signal) and/or the different stoichiometry of anti-CD20 antibodies needed for each assay. *MS4A1* KO cells did not trigger a luciferase signal. The same trend was observed in the ADCP assay (Figure 5D). We observed a decrease in the percentage of THP-1 cells associated with RAJI cell with *MS4A1* KO cells, and to lower extent with *IRF8* KO for the three antibodies.

Altogether, these results show that *IRF8* KO confers a marked resistance to RTX, OFA and OBZ in the three Fc-dependent antibody functions tested. The three antibodies displayed similar requirement for IRF8 and effector function profiles.

### CD20 and IRF8 levels correlate in subsets of diffuse large B-cell lymphomas

To further determine how IRF8 may control CD20 expression, we took advantage of a public dataset of patient cancer cell RNA levels and their associated therapeutic outcomes (47). We examined expression of IRF8 and CD20 in 716 DLBCLs present in this database. The expression of CD20 was slightly but significantly higher in germinal-center B cells (GCB) DLBCLs than in activated B-cell (ABC) and unclassified DLBCLs (median of 10.6 [95% CI: 10.4-10.7] for GCB, 10.1 [9.9-10.2] for ABC and 9.9 [9.5-10.3] for unclassified, p<0.0001, Figure 6A). A similar pattern was observed for IRF8 expression with a median of 7.96 [95% CI: 7.8-8.1] for GCB, 7.5[7.4-7.6] for ABC and 7.6[7.5-7.8] for unclassified (p<0.0001, p=0.0002, Figure 6B). We further observed a significant positive correlation between IRF8 and CD20 expression, with a Spearman coefficient of 0.27 for GCB cells and of 0.4 for unclassified lymphomas (Figure 6C). This was not the case for ABC DLBCLs (Figure 6C).

**Figure 6.**
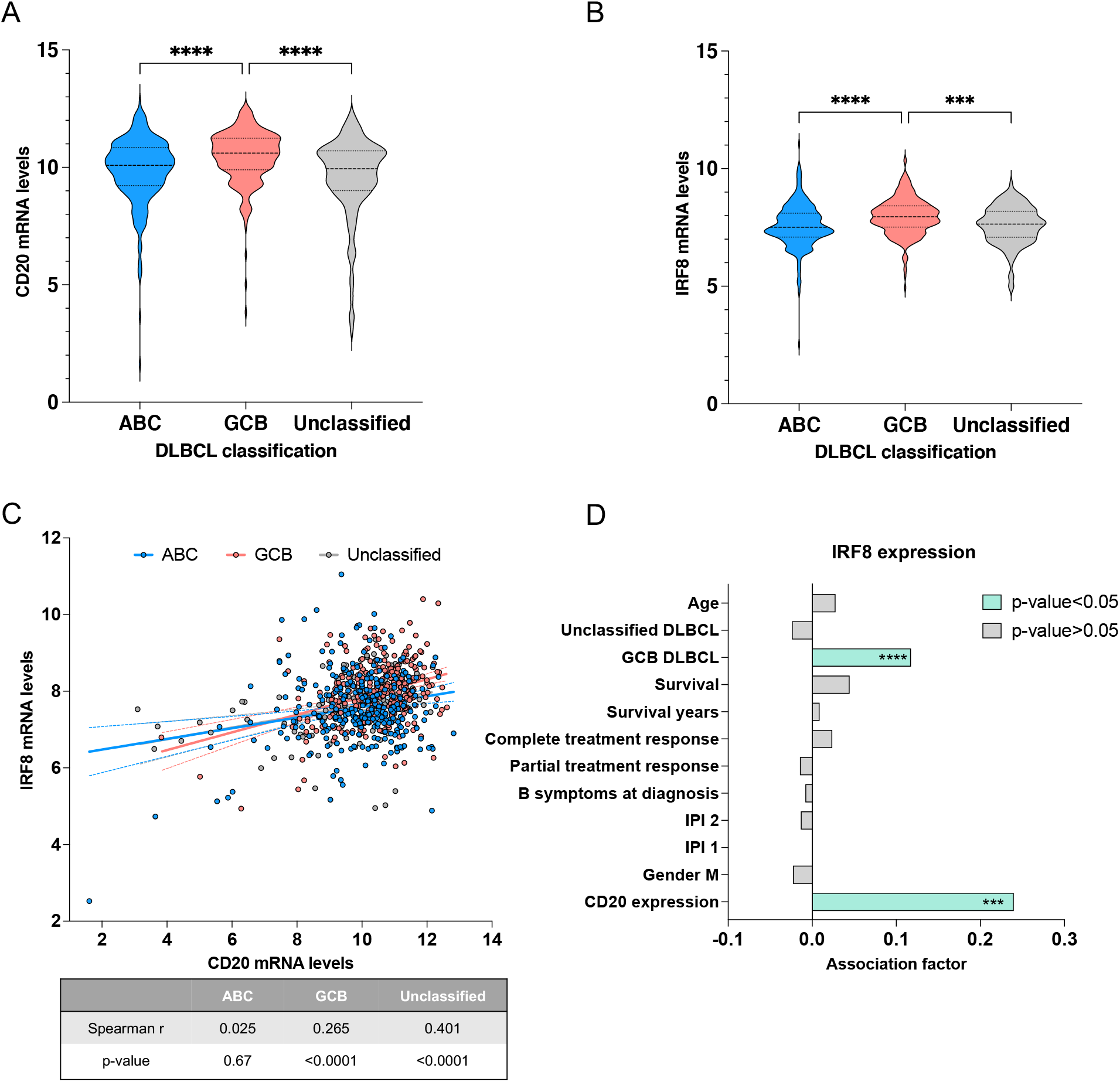
CD20 and IRF8 mRNA levels in diffuse large B-cell lymphomas from 716 patients. **A**. CD20 mRNA levels in DLBCL subsets. Activated B-cell (ABC) subset is in blue, germinal center B-cell (GCB) subset in salmon and unclassified subset in grey. The median is represented in dashed line and quartiles in dotted lines. A Kruskal-Wallis test was performed since normality was not confirmed. ****: p-value <0.0001. **B**. IRF8 mRNA levels in DLBCL subsets. The median is represented in dashed line and quartiles in dotted lines. A Kruskal-Wallis test was performed since normality was not confirmed. ***: p-value = 0.0002, ****: p-value <0.0001. **C**. Correlation between CD20 and IRF8 mRNA levels in DLBCL subsets. Correlation coefficients of Spearman test (r) and p-value are given in the table underneath. Linear regressions are depicted on the figure with 95% confidence intervals. **D**. Parameters associated with IRF8 expression analyzed with the partial least squares (PLS) regression method. The association factor is the estimate effect of each standardized parameter on the standardized CD20 expression taking all parameters into account. Negative and positive factors denote respectively negative and positive associations. Significant associations, as assessed with Jackknife approximate t tests, are in green and ***: p-value<0.0003, ****: p-value<10^−6^.

We also investigated in this dataset the parameters associated with IRF8 expression by performing a multivariate analysis using the partial least regression method. Considering potential cofounder variables such as International Prognostic Indexes, age or treatment response, we observed a strong positive association between IRF8 and CD20 expression (p-value= 1.2·10^−4^) (Figure 6D). GCB DLBCLs were associated with higher IRF8 expression (Figure 6D). No other variables were independently associated to IRF8 expression in this dataset.

Altogether, these results show that IRF8 and CD20 expression levels are associated in GCB and unclassified DLBCL.

## Discussion

We performed a whole-genome CRISPR/Cas9 screening strategy to identify genes involved in the susceptibility of B cells to anti-CD20-induced complement-dependent cytotoxicity. Using two experimental conditions, we described a significant enrichment of gRNAs targeting *IRF8, MS4A1, CUL3, AMBRA1* and *KHDC4* genes in cells surviving CDC. We further knocked-out each individual gene and confirmed a role for *MS4A1* (the gene coding for CD20) and *IRF8* in anti-CD20 susceptibility.

We show that IRF8 is necessary for CD20 expression in RAJI and SU-DHL-4 cells and acts at the transcriptional level. Knocking-down IRF8 decreased CD20 mRNA levels and hence protein production and localization at the cell surface. This is consistent with the presence of an IRF8-binding site in the promoter of *MS4A1* (48). One previous study incidentally reported a down-regulation of CD20 at the surface of RAJI cells treated with IRF8-shRNA without further investigation (49). Our results confirm and extend this observation. Little is known about the transcriptional regulation of CD20 expression. IRF8 is expressed in a variety of non B-cells, including monocytes, macrophages and dendritic cells which do not express CD20 (50). IRF8 is known to form dimers with IRF1, IRF2, AP-1 and PU.1 (51). It is thus likely that other transcription factors, acting in direct or indirect association with IRF8, are necessary to promote CD20 transcription in B cells. Alternatively, the *MS4A1* gene may not be accessible to transcription in non-B cells. We did not detect an upregulation of CD20 after type-I interferon treatment of primary B cells or B cell lines even though there was an upregulation of IRF8 (not shown). Future work is warranted to characterize how IRF8 may be modulated in B cells and how this transcription factor acts to promote CD20 expression.

We observed a correlation between IRF8 and CD20 mRNA levels using RNAseq data from 66 B-cell lines, with differences depending on the type of lymphoma. The correlation was significant for diffuse large B-cell lymphoma, acute lymphocytic leukemia and other non-Hodgkin lymphomas but not for Hodgkin or Burkitt lymphomas. However, Burkitt lymphomas were mostly double positive, and Hodgkin lymphoma lacked CD20 expression. It is likely that the developmental stage of the B cells at the origin of the cancer may impact the role of IRF8 in controlling CD20 expression. Accordingly, we show that primary B cells express at most stages both IRF8 and CD20. We did not detect CD20^+^ B cells lacking IRF8, but we could identify a fraction of tissue-like and activated memory B cells that expressed IRF8 but not CD20. This confirms that IRF8 is necessary but not sufficient to trigger CD20 transcription. We also investigated the association between CD20 and IRF8 at the RNA level in cells from patients with DLBCL, using published RNAseq data (47). CD20 and IRF8 expressions were higher in GCB DLBCL than in ABC or unclassified DLBCL. There was a significant correlation in the expression of the two genes in GCB and unclassified lymphomas. A multivariate analysis confirmed a positive association of IRF8 expression and GCB subtype with CD20 expression. Altogether, these analyses strongly suggest that IRF8 may control CD20 expression in some, but not all subsets of normal or tumoral B cells.

We further show that knocking-out *IRF8* decreased sensitivity of target cells to Fc-mediated functions of three widely used therapeutic anti-CD20 antibodies, RTX, OFA and OBZ. This decrease was observed in three different cell-based assays: CDC, and surrogates of ADCC and ADCP. The extent of protection in RAJI cells lacking *IRF8* varied with the assay. The highest effect was seen for CDC with 4.1 to 7.7-fold decreased killing, with the three antibodies compared to control cells whereas the decrease of antibody efficacy was between 1.4- to 2-fold with the two other assays. This can be explained by the fact that hexamers of antibodies are needed for optimal complement activation (52). The amount of CD20 at the surface has thus to be sufficiently high to enable the complex organization of multiple Fc domains, whereas dimers of antibodies may be sufficient to trigger ADCC (53). It is also likely that the two surrogate assays are more sensitive than authentic ADCC and ACDP events. The effective concentrations 50 (EC50) of the antibodies were for instance 45 to 1000-fold lower in the ADCC assay than in CDC.

What might be the therapeutic consequences of our observation? The regulation of CD20 by IRF8 may play a role in the resistance or relapse to anti-CD20 therapies in use for lymphomas. The *IRF8* gene is known to be a hotspot of mutations. It is found mutated in approximately 10% of DLBCL with mostly missense mutations (47,54), in primary mediastinal large B-cell lymphoma (55) and follicular lymphomas (56). It will be worth determining whether lymphoma cells carrying *IRF8* mutations or downregulating this protein could be selected by anti-CD20 treatment. IRF8 has recently been shown to be a transcriptional regulator of *CD37* in DLBCLs (57) and CD37 loss is associated with CD20 downregulation and predicts worse survival (58). Anti-CD20 antibodies are also widely used to treat autoimmune diseases such as rheumatoid arthritis (RA), multiple sclerosis (MS) and systemic lupus erythematosus (SLE) (59). RA and SLE are interferonopathies (60). Patients with MS are sometimes treated with IFN-β (61). The impact of IFN on expression of IRF8 and CD20 is not well characterized *in vivo*, even if CD20 is not considered as being an ISG. Understanding the role of IRF8, IFN and other cytokines on CD20 expression will require further investigation.

Altogether, our data shed light on a mechanism that regulate CD20 expression in B cells and their sensitivity to anti-CD20 depleting antibodies. Given the importance of CD20 as a therapeutic target, our results may help understanding treatment resistance in patients suffering from cancer and autoimmune diseases.

## Methods

### Cells

RAJI (ATCC® CCL-86™), SU-DHL-4 (ATCC® CRL-2957™), THP-1 (ATCC® TIB-202^™^), Jurkat-CD16-NFAT-rLuc cells (Promega ADCC Reporter Bioassay(#G7102)), 293T cells (ATCC® CRL-3216™) and peripheral blood mononuclear cells (PBMCs) were grown as described (supplemental method).

### Recombinant antibody production

Rituximab, Ofatumumab and Obinutuzumab antibodies were produced as described ^37,38^ (supplemental method). For binding experiments, antibodies were biotinylated using the EZ-Link Micro NHS-PEG_4_-Biotinylation Kit (#21955 ThermoScientific).

### CRISPR-Cas9 genome-wide screen

The genome-wide library Toronto KnockOut (TKO) CRISPR Library – Version 3 (TKOv3, #90294, Addgene) ^39^, was cloned in the lentiviral vector pLentiCRISPRv2 (#52961, Addgene). Lentivectors were produced using calcium phosphate transfection and production titer was determined by ELISA p24. Cells were transduced by spinoculation with 240 ng vector/million cells with 20 μg/mL DEAE-dextran. Transduced RAJI cells were subjected to 2 μg/mL of Rituximab and 35% of normal human serum (NHS). Conditions A and B are detailed in supplemental methods. DNA was extracted using two phenol-chloroform extractions and purified by ethanol precipitation. A nested PCR was performed with a first reaction of 12 cycles and a second of 20 cycles adding a barcode for sequencing. Primers and protocols are detailed in supplemental methods. High-throughput sequencing was performed with NextSeq 500 (#20024906 Illumina). gRNA enrichment was then analyzed using Model-based Analysis of Genome-wide CRISPR-Cas9 Knockout (MAGeCK) ^40,41^.

### Individual knock-out cells

Transient knock-out cells were generated by electroporation using Lonza 4D-Nucleofector® X-Unit with ribonucleoproteins. More details are in supplemental methods.

### Assessment of individual KO cells

DNA was extracted from KO cells using the QIAamp® DNA Mini Kit (#51304, Qiagen), followed by a genomic PCR with Q5® High-Fidelity DNA Polymerase (#M0491S, New England BioLabs) kit. PCR products were sequenced. KO efficiency was assessed using Tracking of Indels by Decomposition (TIDE)^42^.

### Flow cytometry

Cells were stained in MACS Buffer (PBS + 1.25 g/L Bovine Serum Albumine (BSA) + 2 mM EDTA) for surface staining and in PBS/BSA 1%/Azide for intracellular staining. Cells (1×10^5^) were incubated with antibodies listed in supplemental methods for 30 min at 4°C. Biotinylated antibodies were incubated for 30 min at 4°C in MACS Buffer. Alexa Fluor™ 647 conjugated streptavidin (#S21374, ThermoFisher Scientific) was then added and incubated for 30 min at 4°C. Cells were permeabilized with 1% triton and incubated with anti-IRF8 antibody. Data were acquired with an Attune NxT instrument (Life Technologies).

### Fc-dependent functions assays

For flow cytometry CDC assay, cells (1×10^5^) were incubated with RTX (10 μg/mL unless otherwise stated). Human normal serum was added as a source of complement at the indicated dilutions. Cells were stained with LIVE/DEAD Fixable Aqua Dead Cell marker. Data were acquired on an Attune NxT instrument (Life Technologies) and CDC was calculated as 100 × (% of dead cells with serum − % of dead cells without serum)/(100 − % of dead cells without serum). For high-throughput microscopy, 1×10^5^ RAJI cells were seeded in a 96-well flat bottom black plate (#655090, Greiner Bio-One) 12 μg/mL RTX, 25% NHS, 1.25×10^−1^ μg/mL Hoescht and iodide propidium. ADCC activity was assessed with ADCC Reporter Bioassay cells from Promega (supplemental methods). ADCP activity was assessed as described in ^43^ and supplemental methods.

### Analysis of RNAseq data

RNAseq data were obtained from the Cancer Cell Line Encyclopedia^44^ available online (https://depmap.org/portal/download/). Gene counts were normalized using upper quartile and logarithm 2. Access to RNA-seq data from DLBCL patients ^45^ was authorized through an access agreement signed with Pr Sandeep Dave’s laboratory and exempted of ethical evaluation by the Institutional Review Board under 45CFR46.104(d)(4)(ii) legislation. The mapped RNA sequencing reads from 775 samples were downloaded from European Genome-Phenome Archive (EGA: EGAS00001002606).

### Statistical analysis

Flow cytometry data were analyzed with FlowJo v10 software (TriStar). Calculations were performed using Excel 365 (Microsoft) and figures were drawn on Prism 9 (GraphPad Software). Partial least square regression with “leave-one-out” strategy was performed with R version 4.0.2 on RStudio Desktop 1.3.959 (R Studio, PBC) using the Package pls version 2.7-3.

## Acknowledgments

We thank members of the Virus & Immunity Unit and Maaran Michael RAJAH for their support. We thank Nicoletta Casartelli for critical reading of the manuscript. We thank the staff of UTechS Cytometry and Biomarkers, C2RT from Institut Pasteur for their technical help and Pascal Campagne from The Center of Bioinformatics, Biostatistics and Integrative Biology of Institut Pasteur for his statistical expertise. Work in OS lab is funded by Institut Pasteur, Urgence COVID-19 Fundraising Campaign of Institut Pasteur, Fondation pour la Recherche Médicale (FRM), ANRS, the Vaccine Research Institute (ANR-10-LABX-77), Labex IBEID (ANR-10-LABX-62-IBEID), ANR/FRM Flash Covid PROTEO-SARS-CoV-2, ANR Coronamito, and IDISCOVR European Health Emergency Preparedness and Response Authority (HERA). LG is supported by the French Ministry of Higher Education, Research and Innovation.

## Author Contributions

Experimental strategy and design: LG, FR, TB, and OS

Funding acquisition: OS, MV, HM

Laboratory Experiments: LG, FR, FP, PHC, CP and HM

Bioinformatics analysis: AV, AB and LG

Manuscript writing: LG

Manuscript editing: LG, TB, FR and OS

All authors reviewed and approved the final version of the manuscript.

## Disclosure of Conflicts of Interest

The authors have no conflict of interest to disclose.

## Notes

### Competing Interest Statement

The authors have declared no competing interest.

## References

1. Stashenko P, Nadler LM, Hardy R, Schlossman SF. Characterization of a human B lymphocyte-specific antigen. The Journal of Immunology. 1 oct 1980;125(4):1678–85.

2. van Lochem E g., van der Velden V h. j., Wind H k., te Marvelde J g. Westerdaal N a. c., van Dongen J j. m. Immunophenotypic differentiation patterns of normal hematopoiesis in human bone marrow: Reference patterns for age-related changes and disease-induced shifts. Cytometry Part B: Clinical Cytometry. 2004;60B(1):1–13.

3. Loken MR, Shah VO, Dattilio KL, Civin CI. Flow Cytometric Analysis of Human Bone Marrow. II. Normal B Lymphocyte Development. Blood. 1 nov 1987;70(5):1316–24.

4. Bubien JK, Zhou LJ, Bell PD, Frizzell RA, Tedder TF. Transfection of the CD20 cell surface molecule into ectopic cell types generates a Ca2+ conductance found constitutively in B lymphocytes. J Cell Biol. juin 1993;121(5):1121–32.

5. Li H, Ayer LM, Lytton J, Deans JP. Store-operated Cation Entry Mediated by CD20 in Membrane Rafts*. Journal of Biological Chemistry. 24 oct 2003;278(43):42427–34.

6. Tedder TF, Boyd AW, Freedman AS, Nadler LM, Schlossman SF. The B cell surface molecule B1 is functionally linked with B cell activation and differentiation. J Immunol. août 1985;135(2):973–9.

7. Plosker GL, Figgitt DP. Rituximab. Drugs. 1 avr 2003;63(8):803–43.

8. Siegel RL, Miller KD, Fuchs HE, Jemal A. Cancer statistics, 2022. CA: A Cancer Journal for Clinicians. 2022;72(1):7–33.

9. Grillo-López AJ, Hedrick E, Rashford M, Benyunes M. Rituximab: Ongoing and future clinical development. Seminars in Oncology. 1 févr 2002;29(1, Supplement 2):105–12.

10. Lee DSW, Rojas OL, Gommerman JL. B cell depletion therapies in autoimmune disease: advances and mechanistic insights. Nat Rev Drug Discov. 2021;20(3):179–99.

11. Niederfellner G, Lammens A, Mundigl O, Georges GJ, Schaefer W, Schwaiger M, et al. Epitope characterization and crystal structure of GA101 provide insights into the molecular basis for type I/II distinction of CD20 antibodies. Blood. 14 juill 2011;118(2):358–67.

12. Kumar A, Planchais C, Fronzes R, Mouquet H, Reyes N. Binding mechanisms of therapeutic antibodies to human CD20. Science. 14 août 2020;369(6505):793–9.

13. Singh V, Gupta D, Almasan A. Development of Novel Anti-Cd20 Monoclonal Antibodies and Modulation in Cd20 Levels on Cell Surface: Looking to Improve Immunotherapy Response. J Cancer Sci Ther. nov 2015;7(11):347–58.

14. Coiffier B. Rituximab therapy in malignant lymphoma. Oncogene. mai 2007;26(25):3603–13.

15. Pfreundschuh M, Trümper L, Österborg A, Pettengell R, Trneny M, Imrie K, et al. CHOP-like chemotherapy plus rituximab versus CHOP-like chemotherapy alone in young patients with good-prognosis diffuse large-B-cell lymphoma: a randomised controlled trial by the MabThera International Trial (MInT) Group. The Lancet Oncology. 1 mai 2006;7(5):379–91.

16. Li Y, Williams ME, Cousar JB, Pawluczkowycz AW, Lindorfer MA, Taylor RP. Rituximab-CD20 complexes are shaved from Z138 mantle cell lymphoma cells in intravenous and subcutaneous SCID mouse models. J Immunol. 15 sept 2007;179(6):4263–71.

17. Beers SA, French RR, Chan HTC, Lim SH, Jarrett TC, Vidal RM, et al. Antigenic modulation limits the efficacy of anti-CD20 antibodies: implications for antibody selection. Blood. 24 juin 2010;115(25):5191–201.

18. Davis TA, Czerwinski DK, Levy R. Therapy of B-Cell Lymphoma with Anti-CD20 Antibodies Can Result in the Loss of CD20 Antigen Expression1. Clinical Cancer Research. 1 mars 1999;5(3):611–5.

19. Horvat M, Kloboves Prevodnik V, Lavrencak J, Jezersek Novakovic B. Predictive significance of the cut-off value of CD20 expression in patients with B-cell lymphoma. Oncology Reports. 1 oct 2010;24(4):1101–7.

20. Johnson NA, Boyle M, Bashashati A, Leach S, Brooks-Wilson A, Sehn LH, et al. Diffuse large B-cell lymphoma: reduced CD20 expression is associated with an inferior survival. Blood. 16 avr 2009;113(16):3773–80.

21. Hiraga J, Tomita A, Sugimoto T, Shimada K, Ito M, Nakamura S, et al. Down-regulation of CD20 expression in B-cell lymphoma cells after treatment with rituximab-containing combination chemotherapies: its prevalence and clinical significance. Blood. 14 mai 2009;113(20):4885–93.

22. Pierpont TM, Limper CB, Richards KL. Past, Present, and Future of Rituximab—The World’s First Oncology Monoclonal Antibody Therapy. Front Oncol. 4 juin 2018;8:163.

23. Dall’Ozzo S, Tartas S, Paintaud G, Cartron G, Colombat P, Bardos P, et al. Rituximab-dependent cytotoxicity by natural killer cells: influence of FCGR3A polymorphism on the concentration-effect relationship. Cancer Res. 1 juill 2004;64(13):4664–9.

24. Decaup E, Rossi C, Gravelle P, Laurent C, Bordenave J, Tosolini M, et al. A Tridimensional Model for NK Cell-Mediated ADCC of Follicular Lymphoma. Front Immunol. 2019;10:1943.

25. Minard-Colin V, Xiu Y, Poe JC, Horikawa M, Magro CM, Hamaguchi Y, et al. Lymphoma depletion during CD20 immunotherapy in mice is mediated by macrophage FcγRI. FcγRIII, and FcγRIV. Blood. 15 août 2008;112(4):1205–13.

26. Uchida J, Hamaguchi Y, Oliver JA, Ravetch JV, Poe JC, Haas KM, et al. The Innate Mononuclear Phagocyte Network Depletes B Lymphocytes through Fc Receptor–dependent Mechanisms during Anti-CD20 Antibody Immunotherapy. J Exp Med. 21 juin 2004;199(12):1659–69.

27. Carlotti E, Palumbo GA, Oldani E, Tibullo D, Salmoiraghi S, Rossi A, et al. FcγRIIIA and FcγRIIA polymorphisms do not predict clinical outcome of follicular non-Hodgkin’s lymphoma patients treated with sequential CHOP and rituximab. Haematologica. 2007;92(8):1127–30.

28. Ghesquières H, Cartron G, Seymour JF, Delfau-Larue MH, Offner F, Soubeyran P, et al. Clinical outcome of patients with follicular lymphoma receiving chemoimmunotherapy in the PRIMA study is not affected by FCGR3A and FCGR2A polymorphisms. Blood. 27 sept 2012;120(13):2650–7.

29. Persky DO, Dornan D, Goldman BH, Braziel RM, Fisher RI, LeBlanc M, et al. Fc gamma receptor 3a genotype predicts overall survival in follicular lymphoma patients treated on SWOG trials with combined monoclonal antibody plus chemotherapy but not chemotherapy alone. Haematologica. juin 2012;97(6):937–42.

30. Kennedy AD, Beum PV, Solga MD, DiLillo DJ, Lindorfer MA, Hess CE, et al. Rituximab Infusion Promotes Rapid Complement Depletion and Acute CD20 Loss in Chronic Lymphocytic Leukemia. The Journal of Immunology. 1 mars 2004;172(5):3280–8.

31. Jin X, Ding H, Ding N, Fu Z, Song Y, Zhu J. Homozygous A polymorphism of the complement C1qA276correlates with prolonged overall survival in patients with diffuse large B cell lymphoma treated with R-CHOP. Journal of Hematology & Oncology. 16 août 2012;5(1):51.

32. Racila E, Link BK, Weng WK, Witzig TE, Ansell S, Maurer MJ, et al. A Polymorphism in the Complement Component C1qA Correlates with Prolonged Response Following Rituximab Therapy of Follicular Lymphoma. Clinical Cancer Research. 16 oct 2008;14(20):6697–703.

33. Grandjean CL, Montalvao F, Celli S, Michonneau D, Breart B, Garcia Z, et al. Intravital imaging reveals improved Kupffer cell-mediated phagocytosis as a mode of action of glycoengineered anti-CD20 antibodies. Sci Rep. 4 oct 2016;6:34382.

34. Ivanov A, Beers SA, Walshe CA, Honeychurch J, Alduaij W, Cox KL, et al. Monoclonal antibodies directed to CD20 and HLA-DR can elicit homotypic adhesion followed by lysosome-mediated cell death in human lymphoma and leukemia cells. J Clin Invest. 3 août 2009;119(8):2143–59.

35. Semac I, Palomba C, Kulangara K, Klages N, van Echten-Deckert G, Borisch B, et al. Anti-CD20 Therapeutic Antibody Rituximab Modifies the Functional Organization of Rafts/Microdomains of B Lymphoma Cells1. Cancer Research. 15 janv 2003;63(2):534–40.

36. Weiner GJ. Rituximab: Mechanism of Action. Seminars in Hematology. 1 avr 2010;47(2):115–23.

37. Hayashi K, Nagasaki E, Kan S, Ito M, Kamata Y, Homma S, et al. Gemcitabine enhances rituximab-mediated complement-dependent cytotoxicity to B cell lymphoma by CD20 upregulation. Cancer Sci. mai 2016;107(5):682–9.

38. Mankaï A, Bordron A, Renaudineau Y, Martins-Carvalho C, Takahashi S, Ghedira I, et al. Purine-Rich Box-1–Mediated Reduced Expression of CD20 Alters Rituximab-Induced Lysis of Chronic Lymphocytic Leukemia B Cells. Cancer Research. 14 sept 2008;68(18):7512–9.

39. Pavlasova G, Mraz M. The regulation and function of CD20: an « enigma » of B-cell biology and targeted therapy. Haematologica. juin 2020;105(6):1494–506.

40. Sugimoto T, Tomita A, Hiraga J, Shimada K, Kiyoi H, Kinoshita T, et al. Escape mechanisms from antibody therapy to lymphoma cells: Downregulation of CD20 mRNA by recruitment of the HDAC complex and not by DNA methylation. Biochemical and Biophysical Research Communications. 4 Déc 2009;390(1):48–53.

41. Ushmorov A, Leithäuser F, Sakk O, Weinhaüsel A, Popov SW, Möller P, et al. Epigenetic processes play a major role in B-cell-specific gene silencing in classical Hodgkin lymphoma. Blood. 15 mars 2006;107(6):2493–500.

42. Hart T, Tong AHY, Chan K, Van Leeuwen J, Seetharaman A, Aregger M, et al. Evaluation and Design of Genome-Wide CRISPR/SpCas9 Knockout Screens. G3 (Bethesda). 26 juin 2017;7(8):2719–27.

43. Li W, Xu H, Xiao T, Cong L, Love MI, Zhang F, et al. MAGeCK enables robust identification of essential genes from genome-scale CRISPR/Cas9 knockout screens. Genome Biology. 5 déc 2014;15(12):554.

44. Brinkman EK, Chen T, Amendola M, van Steensel B. Easy quantitative assessment of genome editing by sequence trace decomposition. Nucleic Acids Research. 16 déc 2014;42(22):e168.

45. Schiro K, Worzella T, Cheng ZJ, Garvin D, Surowy T, Hengstl T, et al. Performing ADCC Reporter Bioassay with an environmentally-controlled microplate reader. Drug Discovery World. 1 janv 2014;15:46–8.

46. Miller ML, Finn OJ. Flow cytometry-based assessment of direct-targeting anti-cancer antibody immune effector functions. Methods Enzymol. 2020;632:431–56.

47. Reddy A, Zhang J, Davis NS, Moffitt AB, Love CL, Waldrop A, et al. Genetic and Functional Drivers of Diffuse Large B Cell Lymphoma. Cell. 5 oct 2017;171(2):481–494.e15.

48. Shin DM, Lee CH, Morse HC. IRF8 Governs Expression of Genes Involved in Innate and Adaptive Immunity in Human and Mouse Germinal Center B Cells. PLoS One. 11 nov 2011;6(11):e27384.

49. Słabicki M, Lee KS, Jethwa A, Sellner L, Sacco F, Walther T, et al. Dissection of CD20 regulation in lymphoma using RNAi. Leukemia. déc 2016;30(12):2409–12.

50. Tamura T, Kurotaki D, Koizumi S ichi. Regulation of myelopoiesis by the transcription factor IRF8. Int J Hematol. 1 avr 2015;101(4):342–51.

51. Salem S, Salem D, Gros P. Role of IRF8 in immune cells functions, protection against infections, and susceptibility to inflammatory diseases. Hum Genet. 1 juin 2020;139(6):707–21.

52. Diebolder CA, Beurskens FJ, de Jong RN, Koning RI, Strumane K, Lindorfer MA, et al. Complement is activated by IgG hexamers assembled at the cell surface. Science. 14 mars 2014;343(6176):1260–3.

53. Murin CD. Considerations of Antibody Geometric Constraints on NK Cell Antibody Dependent Cellular Cytotoxicity. Frontiers in Immunology [Internet]. 2020 [cité 14 mars 2022];11. Disponible sur: https://www.frontiersin.org/article/10.3389/fimmu.2020.01635

54. Chapuy B, Stewart C, Dunford AJ, Kim J, Kamburov A, Redd RA, et al. Molecular subtypes of diffuse large B cell lymphoma are associated with distinct pathogenic mechanisms and outcomes. Nat Med. mai 2018;24(5):679–90.

55. Mottok A, Hung SS, Chavez EA, Woolcock B, Telenius A, Chong LC, et al. Integrative genomic analysis identifies key pathogenic mechanisms in primary mediastinal large B-cell lymphoma. Blood. 5 sept 2019;134(10):802–13.

56. Li H, Kaminski MS, Li Y, Yildiz M, Ouillette P, Jones S, et al. Mutations in linker histone genes HIST1H1 B, C, D, and E; OCT2 (POU2F2); IRF8; and ARID1A underlying the pathogenesis of follicular lymphoma. Blood. 6 mars 2014;123(10):1487–98.

57. Elfrink S, ter Beest M, Janssen L, Baltissen MP, Jansen PWTC, Kenyon AN, et al. IRF8 is a transcriptional activator of CD37 expression in diffuse large B-cell lymphoma. Blood Advances. 4 avr 2022;6(7):2254–66.

58. Xu-Monette ZY, Li L, Li L, Byrd JC, Jabbar KJ, Manyam GC, et al. Assessment of CD37 B-cell antigen and cell of origin significantly improves risk prediction in diffuse large B-cell lymphoma. Blood. 29 déc 2016;128(26):3083–100.

59. Du FH, Mills EA, Mao-Draayer Y. Next-generation anti-CD20 monoclonal antibodies in autoimmune disease treatment. Autoimmun Highlights. déc 2017;8(1):1–12.

60. Muskardin TLW, Niewold TB. Type I interferon in rheumatic diseases. Nat Rev Rheumatol. 21 mars 2018;14(4):214–28.

61. Hauser SL, Cree BAC. Treatment of Multiple Sclerosis: A Review. Am J Med. déc 2020;133(12):1380–1390.e2.

62. Tiller T, Meffre E, Yurasov S, Tsuiji M, Nussenzweig MC, Wardemann H. Efficient generation of monoclonal antibodies from single human B cells by single cell RT-PCR and expression vector cloning. Journal of Immunological Methods. 1 janv 2008;329(1):112–24.

63. Wang B, Wang M, Zhang W, Xiao T, Chen CH, Wu A, et al. Integrative analysis of pooled CRISPR genetic screens using MAGeCKFlute. Nat Protoc. mars 2019;14(3):756–80.

64. Barretina J, Caponigro G, Stransky N, Venkatesan K, Margolin AA, Kim S, et al. The Cancer Cell Line Encyclopedia enables predictive modelling of anticancer drug sensitivity. Nature. mars 2012;483(7391):603–7.

65. Liao Y, Smyth GK, Shi W. featureCounts: an efficient general purpose program for assigning sequence reads to genomic features. Bioinformatics. 1 avr 2014;30(7):923–30.

66. Ritchie ME, Phipson B, Wu D, Hu Y, Law CW, Shi W, et al. limma powers differential expression analyses for RNA-sequencing and microarray studies. Nucleic Acids Res. 20 avr 2015;43(7):e47.

