## Supplemental figures and methods for "IRF8 regulates efficacy of therapeutic anti-CD20 monoclonal antibodies"

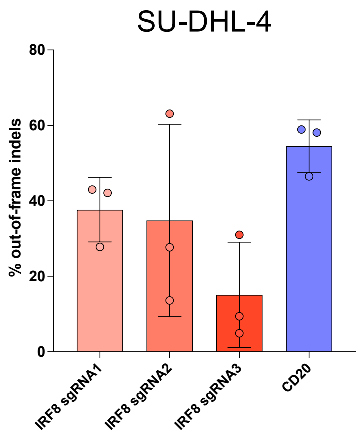


**Figure S1. Characterization of individual knockout in SU-DHL-4 cells** Percentage of out-of-frame indels in SU-DHL-4 cells KO for IRF8 with 3 gRNAs or CD20 KO.

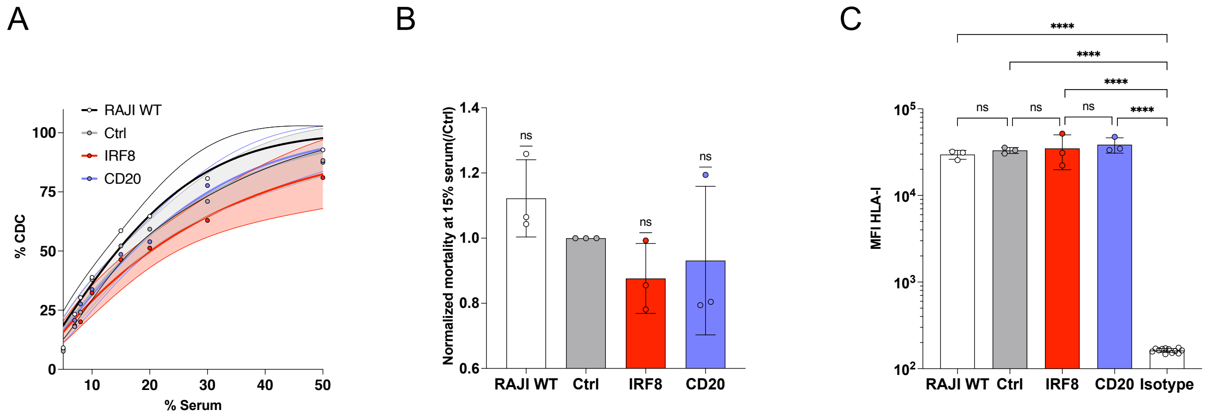


**Figure S2. Susceptibility of KO cells to anti-HLA-I-mediated CDC**

**A.** Percentage of CDC in RAJI WT, Ctrl, IRF8 or CD20 KO. A linear quadratic survival curve with Y as the percentage of dead cells was fitted and the 95% confidence bands were plotted. **B.** Mortality at 15% NHS normalized on mortality of control cells. A Wilcoxon Signed Rank Test was performed. **C.** Median of fluorescence of HLA-I staining in RAJI KO cells. A one-way ANOVA was performed, ****: p-value<0.0001.

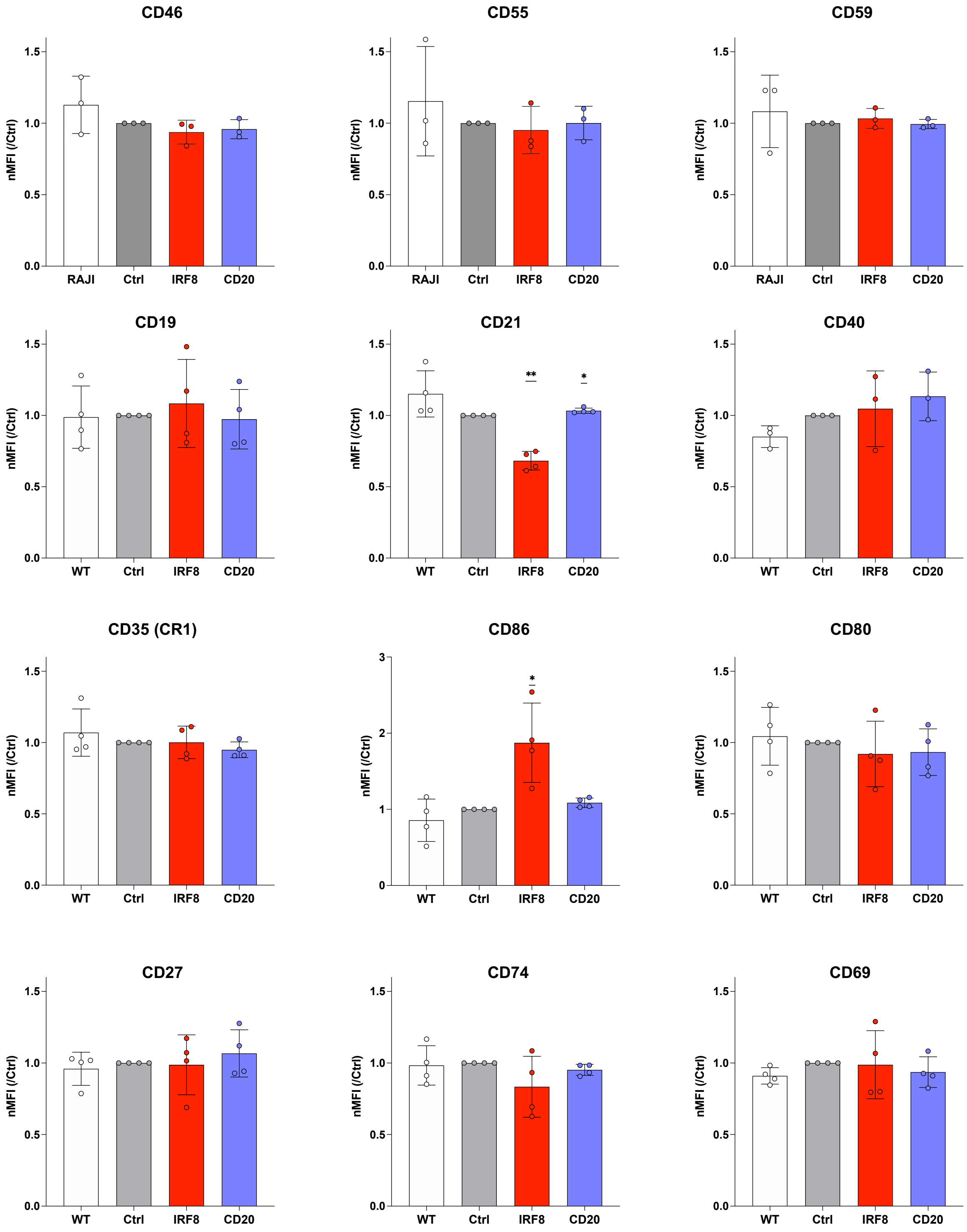


**Figure S3. Impact of KO on several surface protein levels involved in B cell development or CDC resistance**

Expression of the main CDC-regulating proteins (CR1, CD46, CD55 and CD59) and proteins involved in B-cell development and activation expressed as the MFI normalized on control MFI. Differences were assessed with one sample t-tests except for CD59 and CD27 (failed at Shapiro-Wilk test normality test, Wilcoxon signed ranked test). *: $p<0.05$, **: $p<0.005$.

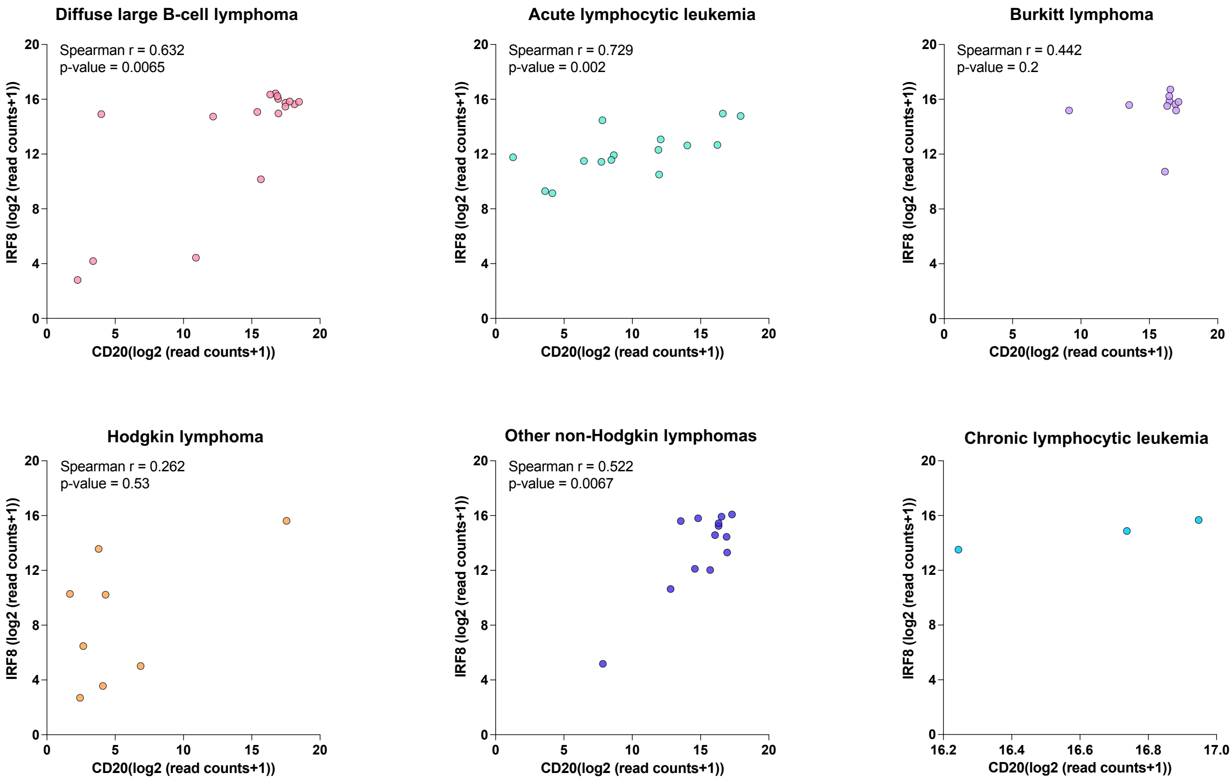


**Figure S4. Correlation between IRF8 and CD20 expression in B cell lines**

66 B cell subsets (Diffuse large B-cell, Hodgkin, Burkitt or other non-Hodgkin lymphomas and acute lymphocytic or chronic lymphocytic leukemias) were analyzed for IRF8 and CD20 mRNA levels. Gene expressions are expressed as the logarithm 2 of read counts+1. Spearman correlation coefficients r and p-values are detailed.

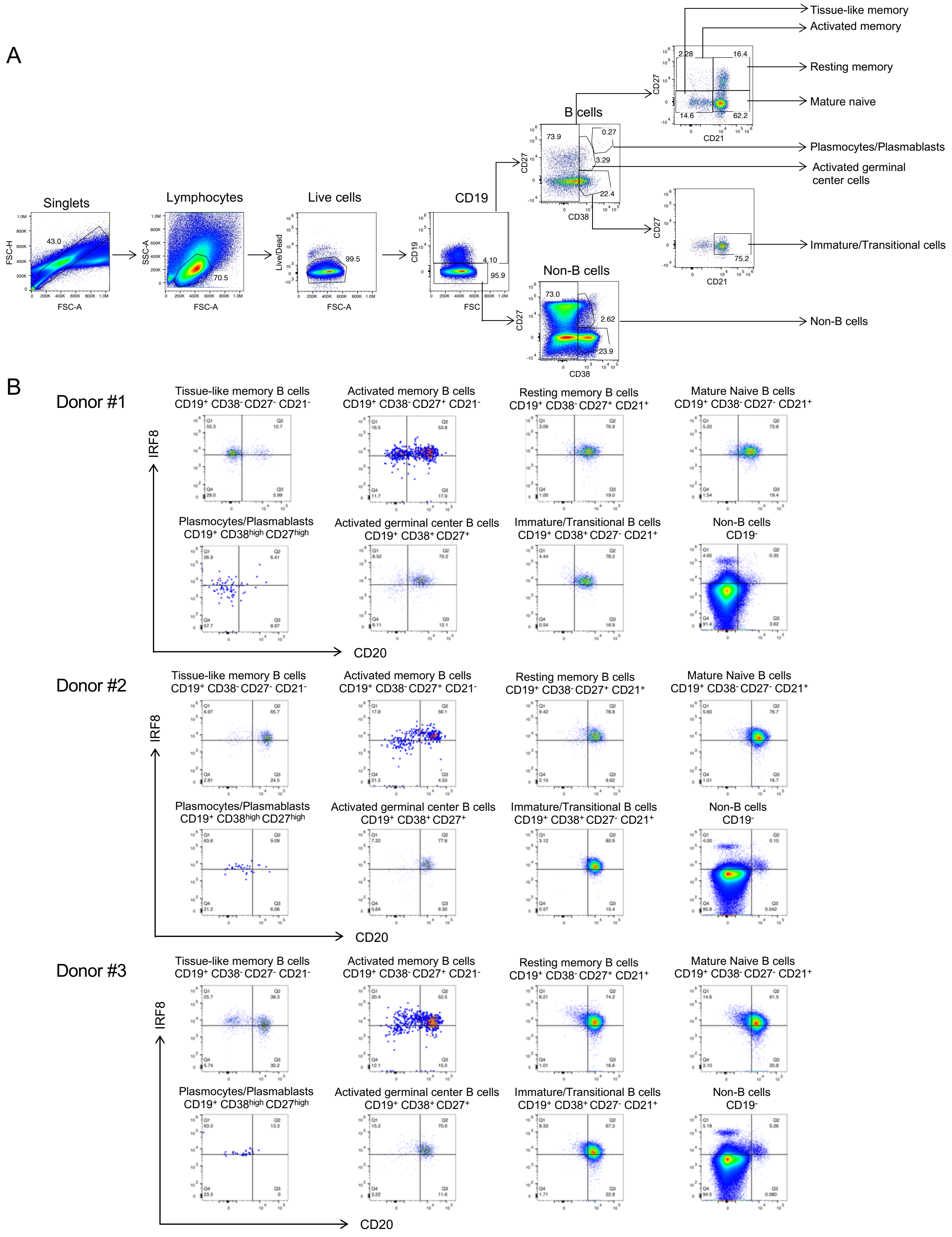


**Figure S5. Stainings of primary B cells with differentiations markers and IRF8**

**A.** Gating strategy used for flow cytometry analysis to identify tissue-like, activated or resting memory, mature naïve, activated germinal center, immature/transitional B cells, plasmocytes/plasmablasts and non-B cells. **B.** IRF8 and CD20 double stainings for each subset of cells in 3 donors.

A

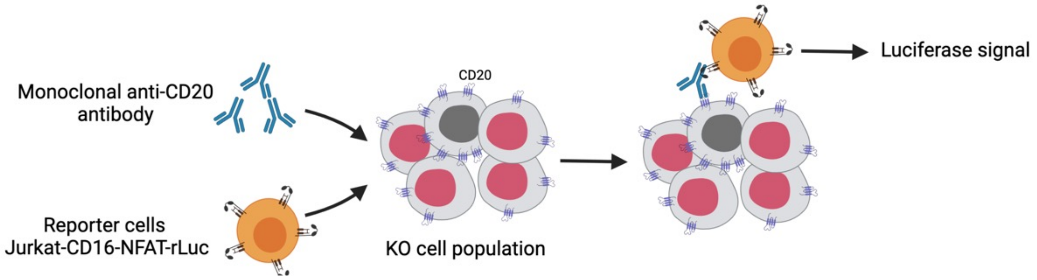


B

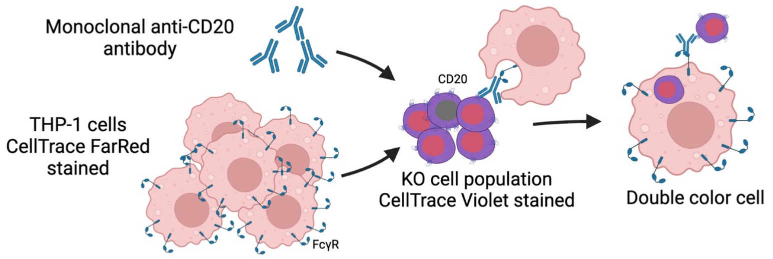


C

| EC50 (SD) | CDC | | | Surrogate ADCC | | | Surrogate ADCP | | |
|---|---|---|---|---|---|---|---|---|---|
| KO \ Ab | RTX | OFA | OBZ | RTX | OFA | OBZ | RTX | OFA | OBZ |
| Ctrl | 1688 (376.8) | 1325 (505.3) | 2476 (1233) | 37.58 (1.992) | 1.255 (0.54) | 4.095 (0.17) | 12.76 (6.02) | 4.338 (1.01) | 6.461 (2.01) |
| IRF8 | 979.4 (405.6) | 266.5 (39.73) | - | 215.2 (128.6) | 2.789 (1.87) | 16.03 (5.08 ) | 25.39 (14.5) | 4.93 (1.0) | 10.2 (4.1) |
| CD20 | - | - | - | - | - | - | 13.51 (4.26) | 4.064 (1.22) | 4.595 (1.5) |

D

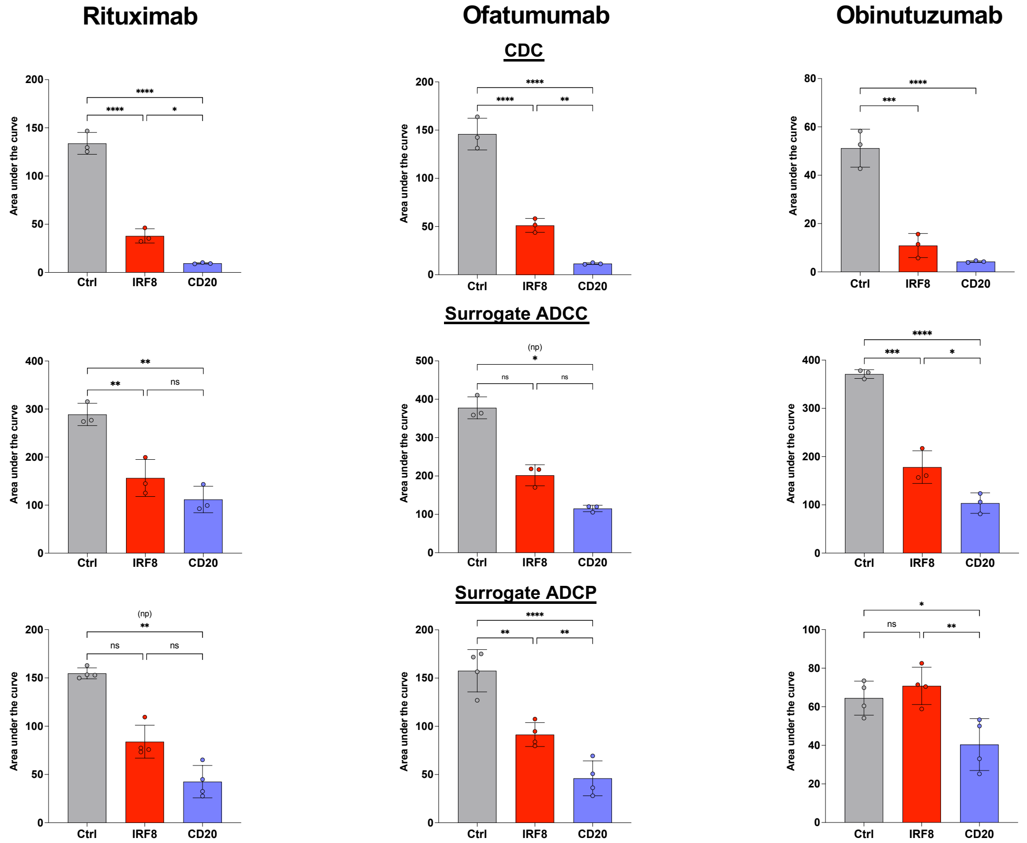


Figure S6. Fc-mediated functions of RTX, OFA and OBZ

**A.** Schematic of the surrogate ADCC assay. KO cells are depicted with a red nucleus. **B.** Schematic of the surrogate ADCP assay with THP-1 and RAJI cells. **C.** Effective concentration 50 of RTX, OFA and OBZ in each test. Standard deviations of 3-4 experiments are in brackets. KO cells were compared to control cells with Mann-Whitney or Kruskal-Wallis tests. Due to the Hook effect, fits for ADCP were calculated for concentrations under 20 ng/mL. **D.** Area under the curve for each curve depicted in figure 5. Differences were assessed with a one-way ANOVA test unless specified by (np) for non-parametric test. *: p-value<0.05, **: p<0.01, ***: p<0.001, ****: p<0.0001

### Supplemental methods

***Cells***

RAJI cells, SU-DHL-4 and THP-1 cells were grown in RPMI medium (#21875034 Gibco) supplemented with 10% Gibco™ Fetal Bovine Serum (FBS) and 1% Penicillin/Streptomycin (PS). Jurkat-CD16-NFAT-rLuc cells, obtained from Promega as ADCC Reporter Bioassay(#G7102) were grown in RPMI with 10% FBS, 1% PS, 1% non-essential amino acids (#11140050 Gibco™), 1mM sodium pyruvate (#11360070 Gibco™), 1 mg/mL G418 (#A2513-5G Clinisciences) and 10 µg/mL hygromycin B (#30240CR Corning). Cells were diluted every 2-3 days to a concentration of $3\text{-}5x10^5$ cells/mL. 293T cells (ATCC® CRL-3216™) were grown in DMEM medium (#10566032 Gibco™) with 10% FBS and 1% PS. Cells were regularly tested negative for mycoplasma contamination (Mycoalert™ Mycoplasma Detection Kit, Lonza). Peripheral blood mononuclear cells (PBMCs) were isolated from healthy human donors from the Établissement Français du Sang following local ethical guidelines.

***Recombinant antibodies production***

Rituximab, Ofatumumab and Obinutuzumab antibodies were produced as previously described (Kumar et al., 2020; Tiller et al., 2008). Briefly, the synthetic codon-optimized DNA fragments coding for the IgH and IgL variable domains of the antibodies were cloned into IgG1-expressing vectors. Antibodies were produced by transient co-transfection of Freestyle™ 293-F suspension cells (ThermoFisher Scientific) using PEI-precipitation method and purified by affinity chromatography (Protein G Sepharose 4 Fast Flow, GE Healthcare). Purified protein fractions were dialyzed against DPBS (Gibco) and digested with PreScission protease.

***CRISPR-Cas9 genome-wide screen***

*Library preparation*

The genome-wide library Toronto KnockOut (TKO) CRISPR Library – Version 3 (TKOv3, #90294, Addgene), described in (Hart et al., 2017), was cloned in the lentiviral vector pLentiCRISPRv2 (#52961, Addgene). To maintain the library complexity, $10^7$ bacterial colonies were kept and DNA was extracted with the QIAGEN Plasmid Giga Kit (#12191, QIAGEN). Lentivectors were produced using calcium phosphate transfection. Briefly, 293T cells were seeded in T175 flasks one day prior to transfection. A DNA precipitate of pLentiCRISPRv2_TKOV3, PAX2 and pVSV-G at a ratio 5:5:2 was formed using calcium chloride solution (#21115 Sigma-Aldrich) and HEPES buffered saline (#51558 Sigma-Aldrich) with a 20–30-minute room temperature incubation. The mix was then added to 293T cells. Lentivectors were harvested at 24, 36 and 48h, filtered through a 0.45 um filter (#16555 Sartorius) and kept at 4°C before use. Supernatants were then pooled and ultracentrifuged at 20,000 *g* for 1h. The production titer was determined by ELISA p24.

*RAJI cells transduction*

$3.6x10^7$ RAJI cells were transduced by spinoculation with 240 ng vector/million cells in a 6-well plate, at 2 million cells/mL with 20 µL/mL DEAE-dextran. The plate was centrifuged at 1100 *g* for 1h at 30°C and 4 mL of complete DMEM was then added. After 48h, cells were selected with 4µg/mL puromycin during 10 days before further use.

*CDC test and fluorescent activated cell sorting*

Transduced RAJI cells (36 million at 1 million cells/mL) were subjected to a complement-dependent cytotoxicity test with 2 µg/mL of Rituximab and 35% of normal human serum (NHS), in duplicate. For condition A, cells were stained with LIVE/DEAD™ Fixable Aqua Dead Cell Stain Kit (ThermoFisher) for 30 min at 4°C and resuspended in PBS + 2% FBS + 25mM HEPES+ 5 mM EDTA after 24h. Live cells were sorted using MoFlo Astrios sorter (Beckman Coulter) at The Cytometry and Biomarkers UTechS platform of Institut Pasteur. As the number of live cells was low, the duplicates were pooled. For condition B, cells were subjected to a CDC test with 1 µg/mL of Rituximab and 50% of normal human serum (NHS) for 24h and then put back in culture in RPMI for 8 days.

*DNA extraction*

Sorted cells were centrifuged 20 min at 500 *g* and lysed with lysis buffer (NaCl 300 mM, SDS 0.1%, EDTA 10mM, EGTA 20 mM and Tris 10 mM supplemented with 3 µg of RNAseA (#R4642-10MG Sigma) and 3 µg of proteinase K (#19133 Qiagen)). After overnight incubation at 65°C, DNA was extracted using two subsequent phenol-chloroform extractions, and purified by ethanol precipitation (overnight incubation at -80°C). DNA was resuspended in water.

*gRNA amplification and high-throughput sequencing*

A nested PCR was performed with a first reaction of 12 cycles and a second of 20 cycles adding a barcode for sequencing. Primers are depicted in the table below.

| | |
|---|---|
| *Reaction 1_ Forward primer* | |
| **PLC-Seq_R1_F** | gagggcctatttcccatgattccttca |
| *Reaction 1_Reverse primer* | |
| **PLC-Seq_R1_R** | aacttctcggggactgtgg |
| *Reaction 2_Forward primer* | |
| **PLC-Seq_R2_TruSeq_1** | AATGATACGGCGACCACCGAGATCTACACTCTTTCCCTACACGACGCTCTTCCGATCTATCACGtcttgtggaaaggacgaaacaccg |
| **PLC-Seq_R2_TruSeq_2** | AATGATACGGCGACCACCGAGATCTACACTCTTTCCCTACACGACGCTCTTCCGATCTCGATGTtcttgtggaaaggacgaaacaccg |
| **PLC-Seq_R2_TruSeq_3** | AATGATACGGCGACCACCGAGATCTACACTCTTTCCCTACACGACGCTCTTCCGATCTTTAGGCtcttgtggaaaggacgaaacaccg |
| *Reaction 2_ Reverse primer* | |
| **PLC-Seq_R2_TruSeq_RS** | CAAGCAGAAGACGGCATACGAGATGTGACTGGAGTTCAGACGTGTGCTCTTCCGATCTtgccactttttcaagttgataacggact |

Small letters: lentivector regions, capital letters: region necessary for sequencing, red letters: barcode

A first PCR was performed with Herculase II Fusion DNA Polymerase kit (#600679, Agilent), 100 mM dNTPs, 100 µM primers and 5% DMSO. DNA was next purified using QIAquick® PCR Purification Kit (Qiagen). A second PCR was performed with different primers for each sample and PCR products were purified with AMPure magnetic beads (# A63880, Beckman Coulter). Samples were equimolarly mixed and 40% of DNA from ϕX bacteriophage was added. High-throughput sequencing was performed with NextSeq 500 (Illumina), NextSeq 500/550 High Output Kit v2.5 (75 Cycles) (#20024906 Illumina).

*gRNA enrichment analysis*

Fastq files from sequencing were deconvoluted to retrieve counts for each sample analyzed and reads were mapped on the human genome. Counts were filtered using a cutoff of 30 reads in control condition. gRNA enrichment was then analyzed using Model-based Analysis of Genome-wide CRISPR-Cas9 Knockout (MAGeCK) (Li et al., 2014b; Wang et al., 2019).

***Individual knock-out cells***

Transient knock-out cells were generated by electroporation using Lonza 4D-Nucleofector® X-Unit with ribonucleoproteins. First, Alt-R CRISPR-Cas9 crRNA for desired gene, Alt-R Cas9 Negative Control (#230192486, Integrated DNA Technologies IDT) and Alt-R® CRISPR-Cas9 tracrRNA (#226577680, IDT) were resuspended at 200 µM in Nuclease-Free Duplex Buffer (#1072570, IDT). Alt-R® Cas9 Electroporation Enhancer (#228869105) was resuspended at 180 µM. TracrRNA-crRNA duplexes were generated by mixing an equal volume of each and incubating at 95°C for 5 min, returning to room temperature with a ramp rate of 0.1°C/s. The duplex formed was mixed with 5 µg of TrueCut™

Cas9 Protein v2 (Invitrogen™), 0.25 µL of duplex if 3 duplexes added or 0.75 µL if only one duplex. During the 10-20 min incubation at RT, the cells were counted and sampled at $6x10^5$ cells per transfection condition, centrifuged. The medium was removed. Cells were resuspended in Nucleofection Buffer (16.4 µL of Nucleofector Solution + 3.6 µL of supplement per condition). The nucleofection buffer used depended on the cell line used: Amaxa SE Cell Line 4D-Nucleofector™ X Kit S (#V4XC-1032, Lonza) for RAJI cells and SF Cell Line 4D-Nucleofector™ X Kit S (#V4XC-2032, Lonza) for SU-DHL-4 cells. In a round bottom 96-well plate, 1.75 µL of RNP + 0.5µL of Electroporation Enhancer followed by 20 µL of cells were added. This mix was then introduced in the nucleofection cuvette and nucleofected with program DS-104 for RAJI cells and program DS-130 for SU-DHL-4. Cells were finally transferred in medium in a 96-well plate and left at 37°C, 5% $CO_2$ before further use.

***Assessment of individual KO cells***

KO cells were pelleted at day 4 and 6 post-electroporation for RAJI and SU-DHL-4, respectively. DNA was extracted using the QIAamp® DNA Mini Kit (#51304, Qiagen), followed by a genomic PCR with Q5® High-Fidelity DNA Polymerase (#M0491S, New England BioLabs) kit: 5µL Q5 Reaction Buffer, 2.5 µL 2mM dNTPs, 1.25 µL of forward and reverse primers, 1 µL DNA, 5 µL Q5 High GC Enhancer, 0.25µL Polymerase and up-to 25 µL water. Thermocycler program: 1X 98°C 30s, 40X (98°C 10s, temperature depending on the gene for 30s, 72 °C 1min), 1X 72°C, 2 min.

PCR products were purified with QIAquick PCR Purification Kit (#28104, Qiagen) or PureLink™ Quick PCR Purification Kit (#K310002, Invitrogen) and sent for sequencing at Eurofins Genomics. KO efficiency was assessed using Tracking of Indels by Decomposition (TIDE) (Brinkman et al., 2014).

***Flow cytometry***

Cells were stained in MACS Buffer (PBS + 1.25 g/L Bovine Serum Albumine (BSA) + 2 mM EDTA) for surface staining and in PBS/BSA 1%/Azide for intracellular IRF8 staining. For surface staining, the cells ($1x10^5$) were incubated with the indicated antibodies for 30 min at 4°C, followed by a washing in PBS and fixation with 4% paraformaldehyde (PFA). The antibodies used were: PE Anti-CD20 (Clone 2H7, #556633, BD Biosciences, 1:20), BV421 Anti-CD20 (Clone 2H7, #562873, BD Biosciences, 1:50), FITC Recombinant Anti-Human CD21 (Clone REA 940, #130-115-515, Miltenyi Biotec, 1:50), APC Anti- CD19 (Clone HIB19, #555415, BD Biosciences, 1:20), PE-Cy5 Anti-CD80 (Clone L307.4, #559370, BD Biosciences, 1:20), FITC Anti-CD35 (Clone E11, #555452, BD Biosciences, 1:20), APC Anti-CD27 (Clone O323, #17-0279-73, eBiosciences, 1:20), FITC Anti-CD74 (Clone MB741, #555540, BD Biosciences, 1:20), APC Anti-CD40 (#21270406S, ImmunoTools, 1:50), PE Anti-CD86 (Clone HA5.2B7, #IM2729U, Beckman Coulter, 1:20), PE Anti-CD69 (Clone L78, #341652, BD Biosciences, 1:20), BV421 Anti-CD21 (Clone B-ly4, #562966, BD Biosciences, 1:100), AF700 Anti-CD19 (Clone HIB19,# 561031, BD Biosciences, 1:100), APC Anti-CD38 (Clone HIT2, # 560980, BD

Biosciences, 1:100), PE-CF594 Anti-CD27 (Clone M-T271, # 562324, BD Biosciences, 1:50), BV605 Anti-IgM (Clone G20-127, #562977, BD Biosciences, 1:100), BV786 Anti-IgG (Clone G18-145, #564230, BD Biosciences), FITC Anti-IgA(Clone IS11-8E10, #130-114-001, Miltenyi, 1:100), APC-H7 Anti-IgD (Clone IA6-2, # 561305, BD Biosciences, 1:100), PerCP-Cy5.5 Anti-CD20 (Clone 2H7, # 302326, BioLegend, 1:50), APC Anti-Human HLA-ABC, Clone W6/32, #555555, BD Biosciences, 1:20).

For analysis of RAJI and SU-DHL-4 cells, the indicated biotinylated antibodies were incubated for 30 min at 4°C in MACS Buffer. After a PBS wash, Alexa Fluor™ 647 conjugated streptavidin (#S21374, ThermoFisher Scientific) was added and incubated for 30 min at 4°C. Cells were then washed and fixed. For IRF8 intracellular staining, after fixation with 4% PFA, cells were permeabilized with 1% triton for 10 min, washed with PBS and incubated with anti-IRF8 antibody (APC Recombinant Anti-IRF8 (Clone REA516, #130-108-170, Miltenyi, 1:20) or PE Recombinant Anti-IRF8 (Clone REA516, #130-122-927, Miltenyi, 1:50) for 45min at room temperature. Cells were then washed. Data were acquired with an Attune NxT instrument (Life Technologies).

***Complement-dependent cytotoxicity (CDC) assay***

For flow cytometry, RAJI or SU-DHL-4 cells ($1x10^5$) were seeded in a 96-well round bottom plate with 10 μg/mL of Rituximab unless otherwise stated. Human normal serum (obtained from the Etablissement Français du Sang EFS Rungis) as a source of complement was added at the indicated dilutions. Cells were incubated for 24h except in figure 2 where the incubation was 1h. Cells were then washed twice with PBS, stained with LIVE/DEAD Fixable Aqua Dead Cell marker (dilution 1:1000 in PBS, Life Technologies #L34957) for 30 min à 4°C and finally fixed with 4% PFA. Data were acquired on an Attune NxT instrument (Life Technologies) and CDC was calculated as 100 × (% of dead cells with serum − % of dead cells without serum)/(100 − % of dead cells without serum).

For high-throughput microscopy, $1x10^5$ RAJI cells were seeded in a 96-well flat bottom black plate (#655090, Greiner Bio-One) in 100 μL RPMI with 12 μg/mL RTX, 25% NHS, $1.25 \cdot 10^{-1}$ μg/mL Hoescht and iodide propidium. Cells were then imaged every 2.6 min.

***Surrogate assay of antibody-dependent cellular cytotoxicity (ADCC)***

ADCC activity was assessed with ADCC Reporter Bioassay cells from Promega. $5x10^4$ reporter cells were co-cultured with 500 RAJI KO cells (at days 5,6 and 9 post-KO) in 100 μL in presence of antibody (RTX, OFA or OBI) serially diluted by a factor 5 and starting at 15 μg/mL. After 18h at 37°C, cells were spined and 50 μL were removed and replaced by 50 μL of Bright-Glo™ reagent (#E2620, Promega). The 100 μL were transferred to 96-well flat bottom white plates (#655083, Greiner Cellstar®) and luciferase signal was assessed after 5 min using Perkin Elmer Wallac 1420 Victor2 Microplate reader. For each antibody, the luciferase signal was normalized on the highest signal obtained in each experiment.

***Surrogate assay of antibody-dependent cellular phagocytosis (ADCP)***

ADCP activity was assessed as described in (Miller and Finn, 2020). Briefly, RAJI KO (days 7, 8, 11 and 13 from 3 independent KO) and THP-1 cells were washed with PBS and stained with CellTrace™ Violet and CellTrace™ Far Red proliferation kit respectively. They were then counted and $1x10^5$ RAJI cells were plated in 50 µL of RPMI in a 96-well round bottom plate. We next added 50 µL of antibody diluted serially by 5-fold starting at 10 µg/mL and incubated at room temperature for 15 min. $2x10^4$ THP-1 cells in 100 µL were then added to each well and plates were incubated at 37°C for an hour. After incubation, cells were washed with PBS and fixed with 4% PBS. The percentage of Far Red$^+$ THP-1 cells was determined after acquisition with an Attune NxT cytometer and normalization on the no antibody condition 100 × (% with antibody − % without antibody)/(100 − % without antibody). Technical duplicates were performed for each condition and the mean of the duplicate was used as the value for one experiment.

***Analysis of RNAseq data from cell lines***

RNA sequencing data were obtained from the Cancer Cell Line Encyclopedia (Barretina et al., 2012) available online (https://depmap.org/portal/download/). The gene counts were normalized using upper quartile and logarithm 2 and analysis of cell lines from Acute Lymphocytic Leukemia (ALL), Chronic Lymphocytic Leukemia (CLL), Burkitt lymphoma, Diffuse Large B-Cell Lymphoma (DLBCL), Hodgkin lymphoma or other non-Hodgkin lymphomas was performed on R.

***Analysis of published RNA-seq data from DLBCL patients***

Access to the data published in (Reddy et al., 2017) was authorized through an access agreement signed with Pr Sandeep Dave's laboratory and exempted of further ethical evaluation by the Institutional Review Board under the 45 CFR 46.104(d)(4)(ii) legislation. The mapped RNA sequencing reads from 775 samples were downloaded from European Genome-Phenome Archive (EGA: EGAS00001002606) from the genomic analysis performed by Reddy and colleagues. This dataset involved Diffused Large B cell lymphoma patients who were uniformly treated with standard-of-care therapy (R-CHOP, a combination therapy of anti-CD20 antibody rituximab, cyclophosphamide, hydroxydaunorubicin, vincristine [oncovin], and prednisone). Clinical information was obtained from Reddy et al. supplementary materials, including sex which was also available within the EGA base. We used featureCounts (subread v1.6.1) (Liao et al., 2014) and the GRCh37.p13 annotation from NCBI to obtain the matrix of counts of fragment by genes from downloaded bam files. We filtered gene level raw counts by keeping only genes with more than 1 fragment per million mapped in more than 34 samples. It conserved 90% of the genes (n=17234). Normalized counts were generated by applying a cyclic loess normalization using the voom function of the limma package (v3.46.0)(Ritchie et al., 2015). When including clinical information, we removed 59 samples with discrepancy between the information retrieved from Reddy et al supplementary materials and the one from the metadata in EGA.

***Statistical analysis***

Flow cytometry data were analyzed with FlowJo v10 software (TriStar). Calculations were performed using Excel 365 (Microsoft) and figures were drawn on Prism 9 (GraphPad Software) which was also used to assess statistical significance between different groups with the tests indicated in each figure legend.

Partial least square regression with "leave-one-out" strategy was performed with R version 4.0.2 on RStudio Desktop 1.3.959 (R Studio, PBC) using the Package pls version 2.7-3. Briefly, numerical data were centered and scaled before performing the regression and the number of components was selected with the selectNcomp function. Significance of the variation coefficients was assessed with Jackknife approximate t tests (jack.test). For other R analysis we used the following packages: corrplot (https://github.com/taiyun/corrplot) and readxl (https://CRAN.R-project.org/package=readxl).
